# Multi-omic profiling of breast cancer cells uncovers stress MAPK-associated sensitivity to AKT degradation

**DOI:** 10.1101/2022.10.11.511726

**Authors:** Emily C. Erickson, Inchul You, Grace Perry, Aurelien Dugourd, Katherine A. Donovan, Claire Crafter, Jeffrey W. Johannes, Stuart Williamson, Jennifer I. Moss, Susana Ros, Robert E. Ziegler, Simon T. Barry, Eric S. Fischer, Nathanael S. Gray, Ralitsa R. Madsen, Alex Toker

## Abstract

Over 50% of human tumors display hyperactivation of the serine/threonine kinase AKT. Despite evidence of clinical efficacy, there remains scope to improve upon the therapeutic window of the current generation of AKT inhibitors. Here we report the development of a second-generation AKT degrader, INY-05-040, which outperformed catalytic AKT inhibition with respect to cellular suppression of AKT-driven phenotypes in breast cancer cell lines. A systematic growth inhibition screen across 288 cancer cell lines confirmed a substantially higher potency for INY-05-040 (median GI50_adj_ = 1.1 µM) compared to our first-generation AKT degrader (INY-03-041; median GI50_adj_ = 3.1 µM), with both compounds outperforming catalytic AKT inhibition with GDC-0068 (median GI50_adj_ > 10 µM). Using multi-omic profiling and causal network integration in breast cancer cells, we demonstrate that the enhanced efficacy of INY-05-040 is associated with sustained suppression of AKT signaling, followed by a potent induction of the stress mitogen activated protein kinase (MAPK) c-Jun N-terminal kinase (JNK). Further integration of growth inhibition assays with publicly available transcriptomic, proteomic, and reverse phase protein array (RPPA) measurements established low baseline JNK signaling as a biomarker for breast cancer sensitivity to AKT degradation. Collectively, our study presents a systematic framework for mapping the network-wide signaling effects of therapeutically relevant compounds, and identifies INY-05-040 as a potent pharmacological suppressor of AKT signaling.

## Introduction

The phosphoinositide 3-kinase (PI3K)/AKT network has a fundamental role in the integration of extracellular growth stimuli to regulate cell metabolism, migration, proliferation, and survival^1^. Aberrant activation of this network is widespread in human cancers, particularly those of the female reproductive system^2^. Numerous therapies targeting PI3K/AKT pathway components have been developed and evaluated for their potential as cancer therapeutics, and some have been clinically approved, including the PI3Kα-specific inhibitor alpelisib (PIQRAY®) for ER+/HER2- breast cancer^3^. Because of its central role in mediating PI3K signaling and frequent hyperactivation across cancer types, the serine/threonine protein kinase AKT has become an attractive therapeutic target^4–6^. Several drugs targeting AKT have been developed and evaluated in clinical trials, including ATP-competitive, allosteric, and covalent pan-AKT inhibitors^7–11^. These inhibitors have yet to be approved for the treatment of cancer. Despite promising outcomes in some phase II and ongoing phase III clinical studies^12^, there is scope to improve the therapeutic window of available AKT-targeting compounds. Importantly, conventional AKT inhibitors are largely cytostatic, not cytotoxic, thus failing to eradicate cancer cells as monotherapies. Consequently, there is an unmet need to identify more potent AKT-targeting drugs, in addition to uncovering cellular mechanisms that contribute to the efficacy of AKT inhibition.

Recently, targeted protein degradation using small molecule degraders, also called <u>PRO</u>teolysis <u>TA</u>rgeting <u>C</u>himeras (PROTACs), has emerged as a novel therapeutic modality and as a tool for the chemical depletion of proteins of interest^13–16^. In many cases, PROTACs display increased selectivity over the inhibitors from which they are designed, which presents advantages in limiting off-target toxicities^17^. Targeted protein degradation can also be used as a tool to understand network rewiring dynamics following near-complete and relatively acute depletion of the protein of interest. Potent and selective AKT-targeting PROTACs have been developed, with improved selectivity and potency over parental AKT inhibitors^18–21^.

Here, we report the development of a second-generation AKT degrader, INY-05-040, that selectively and rapidly (<5 h) degrades all three AKT isoforms, and inhibits downstream signaling and cell proliferation across a panel of 288 cancer cell lines. Using a multi-omics approach, combined with computational network modeling and experimental validation, we uncovered several degrader-selective cellular phenotypes in breast cancer cells, including potent activation of the stress mitogen activated protein kinase (MAPK) c-Jun N-terminal kinase 1 (JNK1). Additional breast cancer cell line analyses revealed that a signature of baseline JNK1 activation predicts lower sensitivity to AKT degradation, suggesting a novel biomarker for future therapeutic stratification.

## Results

### INY-05-040 is an improved, second-generation AKT degrader

We previously reported the development of an AKT-targeting degrader INY-03-041, a heterobifunctional degrader consisting of the catalytic AKT inhibitor GDC-0068 chemically linked to the Cereblon (CRBN) recruiter lenalidomide^7^. Despite the potency and selectivity of INY-03- 041, this degrader exhibited relatively slow (12 h) cellular degradation kinetics of all three AKT isoforms^22^. We therefore developed an improved, second-generation AKT degrader, INY-05- 040, consisting of GDC-0068 chemically conjugated to a Von Hippel-Lindau (VHL) ligand via a ten-hydrocarbon linker (**Fig. 1A**). To generate the matched negative control compound INY-05- 040-Neg (**Fig. 1A**), we incorporated a diastereoisomer of the VHL ligand that substantially loses activity towards VHL^23^. The biochemical selectivity of INY-050-040 was comparable to GDC- 0068 across a panel of 468 kinases (**Fig. S1A**). Proteomic analysis in the T lymphoblast cell line MOLT4, chosen due to expression of all three AKT isoforms, confirmed pan-AKT downregulation following 4-h treatment with 250 nM INY-05-040 (**Fig. S1B**).

**Figure 1.**
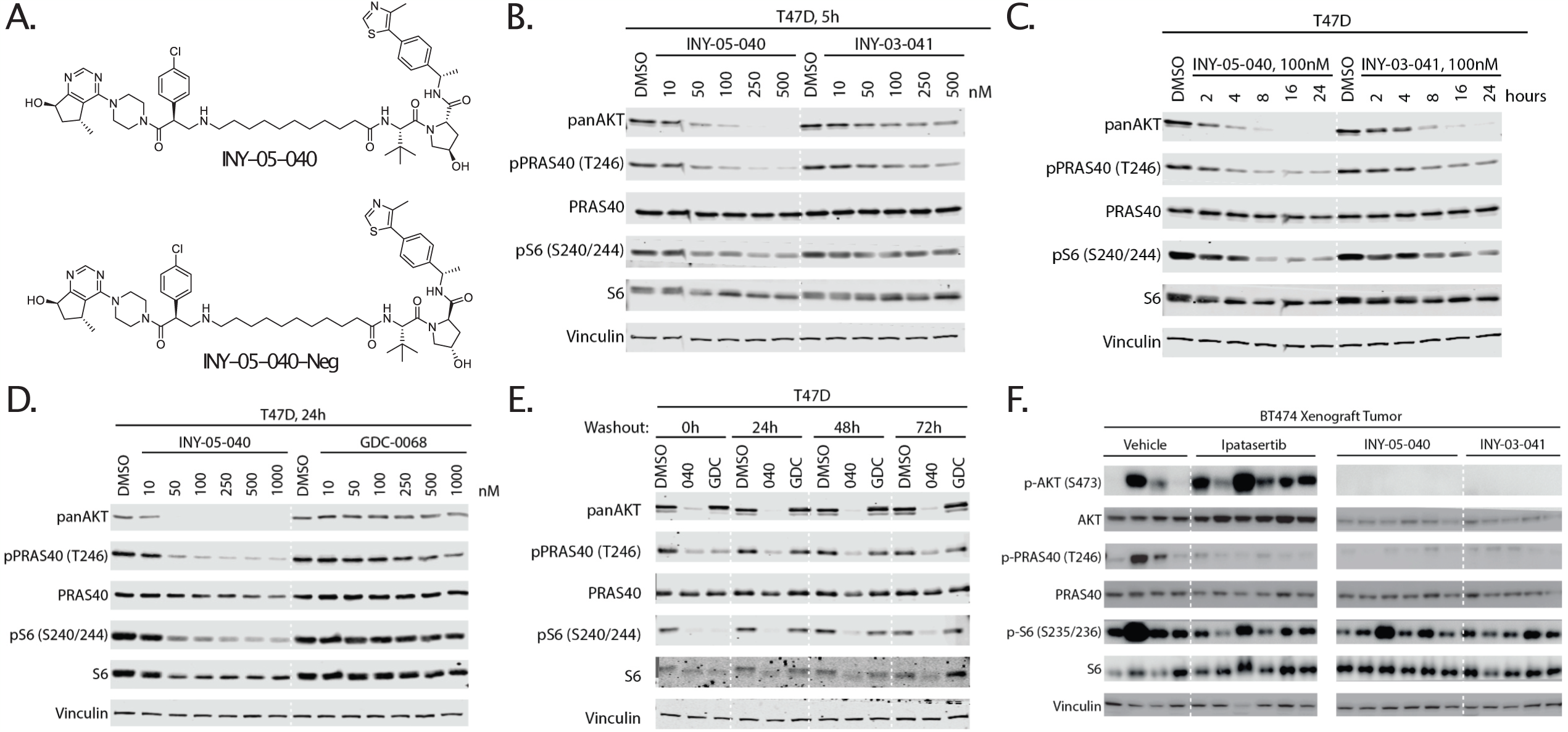
Design and characterization of INY-05-040. (**A**) Chemical structures of INY-05-040 and negative control INY-05-040-Neg. (**B**) Immunoblots for pan-AKT, phospho-PRAS40 (T246), total PRAS40, phospho-S6 (S240/244), total S6, and Vinculin after treatment of T47D cells treated for 5 h with INY-05-040 or INY-03-041 at the indicated concentrations. (**C**) Immunoblots for pan-AKT, phospho-PRAS40 (T246), total PRAS40, phospho-S6 (S240/244), total S6, and Vinculin after treatment of T47D cells treated with INY-05-040 (100 nM) or INY-03-041 (100 nM) for the indicated times. (**D**) Immunoblots for panAKT, phospho-PRAS40 (T246), total PRAS40, phospho-S6 (S240/244), total S6, and Vinculin after 24-h treatment of T47D cells with INY-05- 040 or GDC-0068 at the indicated concentrations. (**E**) Immunoblots for pan-AKT, phospho- PRAS40 (T246), total PRAS40, phospho-S6 (S240/244), total S6, and Vinculin in T47D cells treated with INY-05-040 (100 nM) or GDC-0068 (100 nM) for 5 h followed by washout for the indicated times. (**F**) Immunoblots for pan-AKT, phospho-PRAS40 (T246), total PRAS40, phospho-S6 (S235/236), total S6, and Vinculin in BT-474 mouse xenograft tumors treated with vehicle (10% DMSO, 25% Kleptose), Ipatasertib (12.5 mg/kg), INY-05-040 (25 mg/kg), or INY- 03-041 (25 mg/kg) for 3 days, with a terminal treatment 4 h prior to protein harvest. Additional supporting data related to this figure are included in Fig. S1.

All subsequent evaluation of INY-05-040 was performed in human breast cancer cell lines, due to the high prevalence of PI3K/AKT pathway activation. Exposure of the estrogen receptor-positive (ER+) and *PIK3CA^H^*^1047^*^R^*-mutant T47D cell line to increasing doses of INY-050-040 for 5 h (**Fig.1B**), or over a time course using a dose of 100 nM (**Fig.1C**), revealed a substantially improved dose- and time-dependent reduction in total AKT levels compared to the first-generation degrader, INY-03-041. This was mirrored by potent suppression of downstream PRAS40 (T246) and S6 (S240/S244) phosphorylation (**Fig.1B, 1C**). INY-05-040 also outperformed GDC-0068 in T47D cells treated for 24 h, with >500 nM of GDC-0068 required to achieve comparable signaling suppression to that achieved with 50-100 nM INY-05-040 **(Fig.1D**). As GDC-0068 is also a component of the negative control compound, INY-05-040- Neg, the latter suppressed signaling at higher concentrations (**Fig. S1C**), as expected.

Importantly, unlike non-covalent, catalytic inhibition of AKT with GDC-0068, INY-05-040 treatment of T47D resulted in sustained AKT reduction and suppression of downstream signaling for at least 72 h following compound washout (**Fig. 1E**). Consistent with proteasome-dependent degradation, pharmacological abrogation of proteasomal function or neddylation prevented AKT degradation by INY-05-040 (**Fig. S1F**). We replicated these experiments in the *PTEN*-deficient triple-negative breast cancer (TNBC) cell line MDA-MB-468 (**Fig. S1D, S1E, S1F, S1G**), demonstrating that the favorable cellular properties of INY-05-040 are generalizable across breast cancer cell lineages. Notably, cells exposed to INY-05-040 also exhibited reduced levels of total ribosomal S6 protein, observed within the first 24 h of treatment as well as after compound wash-out (**Fig. 1C, 1D, 1E, S1E**). Consistent with long-term suppression of AKT signaling, both first- and second-generation AKT degraders caused potent suppression of cell growth across four different breast cancer cell lines, at doses well below those required for an equivalent response with catalytic or allosteric AKT inhibitors (**Fig. S1H, S1I)**.

Furthermore, we tested the pharmacodynamic properties of AKT degraders in vivo, using a BT-474C breast cancer xenograft model. Following 4-day-treatment, both first- (INY-03-041) and second-generation (INY-05-040) degraders caused potent reduction in pan-AKT levels, accompanied by downregulation of pPRAS40 (T246) and pS6 (S240/244) (**Fig. 1F**). Likely due to incomplete AKT degradation in vivo, the observed suppression of downstream signaling was equivalent to that observed with GDC-0068.

Taken together, these results show that INY-05-040 is a potent AKT degrader and inhibitor of downstream signaling output, outperforming both our first-generation AKT degrader and GDC-0068.

### Multi-omic profiling reveals AKT degrader-selective responses

To identify mechanisms unique to AKT degradation relative to catalytic inhibition, we performed mRNA sequencing (RNAseq) in T47D breast cancer cells, treated with the respective compounds for 5 or 10 h. To limit the confounding effect of differential potency, we determined the doses of INY-05-040 (100 nM) and GDC-0068 (500 nM) that would result in comparable suppression of downstream signaling at these time points (**Fig. S2**). All studies were performed using nutrient- and growth factor-replete cell culture media.

Consistent with a shared target, the transcriptomes of GDC-0068- and INY-05-040-treated cells clustered closely together, separate from DMSO- and INY-05-040-Neg-treated controls, according to an unsupervised principal component analysis (PCA) (**Fig. 2A**). In agreement with the slower onset of AKT degradation, 5-h treatment with INY-05-040 resulted in differential abundance of only 194 transcripts (100 decreased, 94 increased; absolute fold-change ≥ 1.3), compared to 511 transcripts (249 decreased, 262 increased) with GDC-0068 during the same period (**Fig. 2B**). By contrast, after 10 h, INY-05-040 caused differential abundance of 1394 transcripts (626 decreased, 768 increased; absolute fold-change ≥ 1.3), whereas the extent of GDC-0068-induced transcriptional changes remained stable at 543 transcripts (243 decreased; 300 increased) (**Fig. 2B**). Across all differentially expressed transcripts after 10-h treatment, more than 700 were unique to INY-05-040, compared to less than 100 unique changes for GDC-0068 (**Fig. S3A, S3B**). No differential abundance was observed in response to treatment with the control compound, INY-05-040-Neg (**Fig. 2B**).

**Figure 2.**
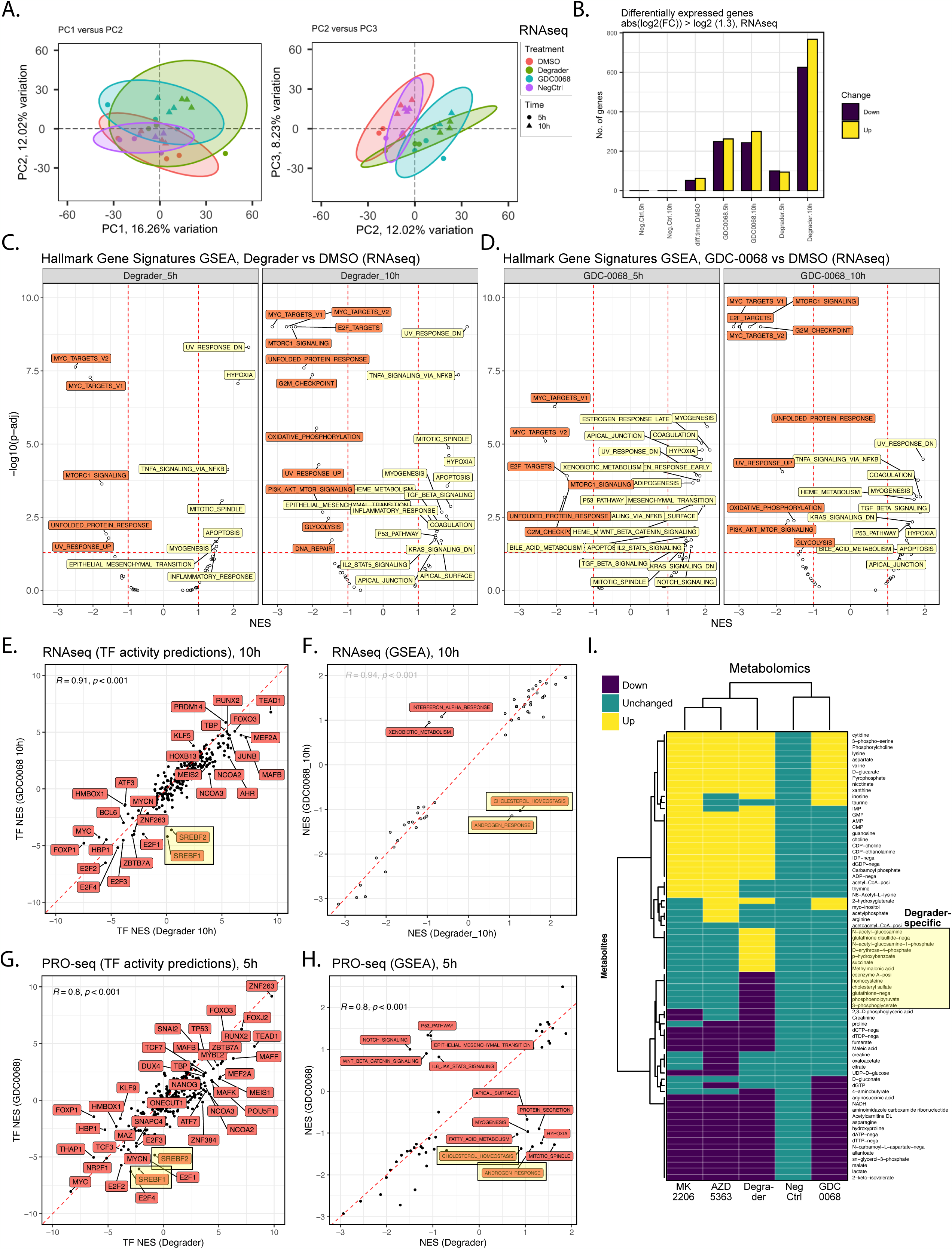
Multi-omic profiling of INY-05-040 and GDC-0068 in T47D breast cancer cells. (**A**) Principal component analysis (PCA) projection of the transcriptomic dataset, comprising n=3 independent experiments per treatment (DMSO; Degrader: 100 nM INY-05-040; 500 nM GDC- 0068; NegCtrl: 100 nM INY-05-040-Neg) and time point (5 h and 10 h). Ellipses are drawn around each group at 95 % confidence level. The first three independent axes (PCs) of highest variation are shown. (**B**) Number of differentially up- and downregulated transcripts (absolute fold-change ≥ 1.3) following differential gene expression analysis (FDR ≤ 0.05) across the indicated comparisons. Comparisons are relative to the corresponding DMSO-treated control; for example, Neg.Ctrl.5h refers to the effect of 10 h treatment with INY-05-040-Neg vs 10 h treatment with DMSO. The exception is “diff.time.DMSO” which evaluates differential expression as a function of time in culture (treatment with DMSO for 10 h versus treatment with DMSO for 5 h). (**C**) + (**D**) Gene set enrichment analysis (GSEA) on the mSigDb HALLMARK collection, based on the ranked *t* values from all genes for the indicated treatments relative to the corresponding DMSO-treated controls. Gene sets are labelled if the absolute normalized enrichment score (NES) exceeds 1 and the adjusted p-value falls below 0.05 (FDR). (**E**) Spearman’s correlation analysis of transcription factor (TF) activity predictions following 10 h treatment with either Degrader or GDC-0068. TF footprint analyses were performed with DoRothEA. SREBF1 (protein name: SREBP1) and SREBF2 (protein name: SREBP2) activity predictions are highlighted due to their divergence between Degrader and GDC-0068, with lower activity predictions observed only in GDC-0068-treated cells. (**F**) Spearman’s correlation analysis of GSEA-derived NES for individual HALLMARK gene sets following 10 h treatment with either Degrader or GDC-0068. “CHOLESTEROL HOMEOSTASIS” and “ANDROGEN RESPONSE” hallmark gene sets are highlighted as having positive and negative NES in Degrader- and GDC-0068-treated cells, respectively. (**G**) As for (E) but based with PRO-seq data corresponding to Degrader and GDC-0068 treatments of T47D cells for 5 h, relative to DMSO-treated control; TF activity predictions were calculated from *t* values from all genes following differential gene expression analysis (FDR ≤ 0.05; n = 2 independent experiments). (**H**) As for (F) but with the PRO-seq data used in (G). (**I**) Hierarchical clustering (Euclidean distance) of differential metabolite abundance (FDR ≤ 0.05) following 24-h treatments of T47D with either AZD 5383 (Capivasertib; catalytic pan-AKT inhibitor; 2 µM), Degrader (INY-05-040; 100 nM), GDC-0068 (catalytic AKT inhibitor; 500-750 nM), MK2205 (allosteric pan-AKT inhibitor; 1 µM) or NegCtrl (INY-05-040-Neg; 100 nM). Differential abundance analysis was performed relative to DMSO-treated controls (n = 9 samples per treatment, from 3 independent experiment with 3 replicates per experiment). More than 85% of the observed differences in metabolite abundance for a given treatment corresponded to at least a 20% change relative to DMSO-treated cells. Metabolite levels that were changed only upon treatment with Degrader are highlighted. Additional supporting data related to this figure are included in Figs. S2, S3, S4.

We next conducted gene set enrichment analysis (GSEA) using the HALLMARK gene signature collection provided by the Broad Institute Molecular Signature Database (MSigDB)^24^. At 10 h, both INY-05-040 and GDC-0068 triggered a transcriptional footprint consistent with suppression of the cell cycle, glycolysis, oxidative phosphorylation, mTORC1 and the unfolded protein response (UPR) (**Fig. 2C, 2D**). Although 5-h treatment with GDC-0068 resulted in a larger number of distinct gene signatures with positive enrichment scores, most of these no longer reached statistical significance after 10 h (**Fig. 2D**), suggesting emerging adaptation to catalytic AKT inhibition. After 10-h treatment, positively enriched gene signatures were largely shared between degrader and catalytic inhibitor, but the underlying gene expression shifts were often more robust following AKT degradation, as evidenced by higher statistical significance despite equivalent sample size (**Fig. 2C, 2D**). Examples include gene signatures related to apoptosis, inflammatory signaling (including TNFα/NFκB) and the mitotic spindle (**Fig. 2C, 2D**).

We next used DoRothEA, a transcriptional footprint-based method featuring a curated gene regulatory network^25^, to predict differences in transcription factor (TF) regulation between INY-05-040 and GDC-0068 at 10 h. Overall, TF activity predictions were highly concordant between the two compounds, with two notable exceptions. The lipid and sterol metabolism-regulating transcription factors, SREBP1 and SREBP2, were predicted as strongly inhibited upon catalytic AKT inhibition but not AKT degradation (**Fig. 2E**). A correlation analysis across the previously generated HALLMARK gene signature enrichments revealed a similar discordance with respect to cholesterol homeostasis and androgen response signatures (**Fig. 2F**). Of note, these two signatures share four transcripts related to lipid and cholesterol synthesis: *SCD, IDI1, HMGCR, HMGCS1*. Both *HMGCR* and *HMGCS1* belong to the list of SREBP1 and SREBP2 targets whose mRNA levels were increased upon 10-h treatment with INY-05-040 but not GDC-0068 (**Fig. S3C**).

These findings were further supported by results from precision nuclear run-on sequencing (PRO-seq), performed after 5-h exposure of T47D cells to INY-05-040 or GDC- 0068. PRO-seq allows mapping of RNA polymerase active sites with base-pair resolution^26^, and changes in the expression of a transcript reflects immediate differences in active transcription, unlike RNAseq which captures steady-state mRNA levels. Similar to the bulk transcriptomes, PRO-seq datasets from degrader- and GDC-0068-treated samples clustered together and away from DMSO-treated controls by PCA (**Fig. S3D**). A substantially higher number of genes were differentially transcribed in response to AKT degradation (**Fig. S3E, S3F**), with further functional enrichment analyses supporting transcriptional regulation of SREBP1/2 and cholesterol homeostasis as defining differences between AKT degradation *versus* catalytic inhibition (**Fig. 2G, 2H**). Such activation of SREBP1 and SREBP2, despite potent AKT/mTORC1 inhibition, would be most consistent with a phenotype of cholesterol depletion^27^.

Given evidence for altered metabolic homeostasis, we next assessed the metabolic profile of T47D cells treated with INY-05-040 and GDC-0068. For comparison, we also included an allosteric (MK-2206) and a second catalytic (AZD 5363) inhibitor. Treatments were performed for 24 h to allow capture of robust and persistent changes, while minimizing the signaling rebound seen with GDC-0068 upon continuous treatment (**Fig. S4**). LC-MS-based metabolomics of 9 cultures per treatment, spanning 3 independent experiments, showed that AKT degradation caused the largest number of differentially abundant metabolites (**Fig. 2I**).

Many metabolite changes were shared across all AKT-targeting compounds, especially MK- 2206 and AZD 5363. Several nucleosides and their phosphorylated derivatives had increased in abundance, including inosine, guanosine, IMP, GMP, AMP and CMP. Metabolite changes unique to treatment with INY-05-040 included intermediates of the hexosamine biosynthesis pathway, the pentose phosphate pathway, glycolysis, the tricarboxylic acid cycle, glutathione and cholesterol metabolism (**Fig. 2I**). Notably, only AKT degradation caused increased levels of methylmalonic acid (MMA), which is known to act as a potent inhibitor of the rate-limiting cholesterol biosynthesis enzyme, HMGCR^28^, and accumulates if vitamin B12 levels are too low relative to the catabolism of branched chain amino acids and odd chain fatty acids^29^.

Taken together, this multi-omic approach supports a widespread perturbation of cellular homeostasis in breast cancer cells treated with INY-05-040, with unique responses to AKT degradation pertaining to cholesterol homeostasis.

### Causal Oriented Search of Multi-Omic Space (COSMOS) identifies altered stress MAPK signaling downstream of AKT degradation

We next reasoned that an integrated, trans-omic integration of the previous datasets may enable us to generate testable mechanistic hypotheses regarding previously unknown signaling changes downstream of AKT degradation. We applied a network analysis approach, COSMOS^30^, to integrate transcriptomic and metabolomic datasets following treatment with AKT degrader INY-05-040 or GDC-0068 for 10 h and 24 h, respectively (**Fig. 3A**). Briefly, COSMOS relies on an extensive prior knowledge network (PKN) of signaling pathways, transcriptional regulation and metabolic reactions, in combination with an Integer Linear Programming (ILP) optimization strategy to identify the smallest coherent subnetwork causally connecting as many deregulated TFs and metabolites in the input data as possible^30, 31^. Input data to COSMOS consisted of the background transcriptome of T47D cells, in addition to treatment-specific DoRothEA-derived TF activity predictions and differentially abundant metabolites. The resulting networks enable identification of top degree signaling nodes, i.e. “hubs”, which are essential for holding a network together due to their high number of connections^32^. Replicate COSMOS runs identified MAPK1 (ERK2) and/or MAPK3 (ERK1) as top degree nodes in both INY-05-040 and GDC-0068 networks (**Fig. 3B, 3C; S5A, S5B**), consistent with the known compensatory RAS/MAPK signaling activation that follows potent PI3K/AKT pathway inhibition^33, 34^. Focusing on unique differences, we noted that the stress MAPKs, MAPK8 (JNK1) and MAPK14 (p38α), were among the top degree nodes in the INY-05-040-specific networks (**Fig. 3B**). MAPK14 was identified as a top degree node in 10 out of 11 COSMOS runs with INY-05-040 input data but was never a top degree node in any of the eight COSMOS runs performed with GDC-0068 input data (**Fig. 3B, 3C**). In two out of eight GDC-0068-specific networks, MAPK14 was not part of the final network; in the remaining six, it featured with a maximum of two connections per network, suggesting a minor role for this kinase in the cellular response to GDC-0068.

**Figure 3.**
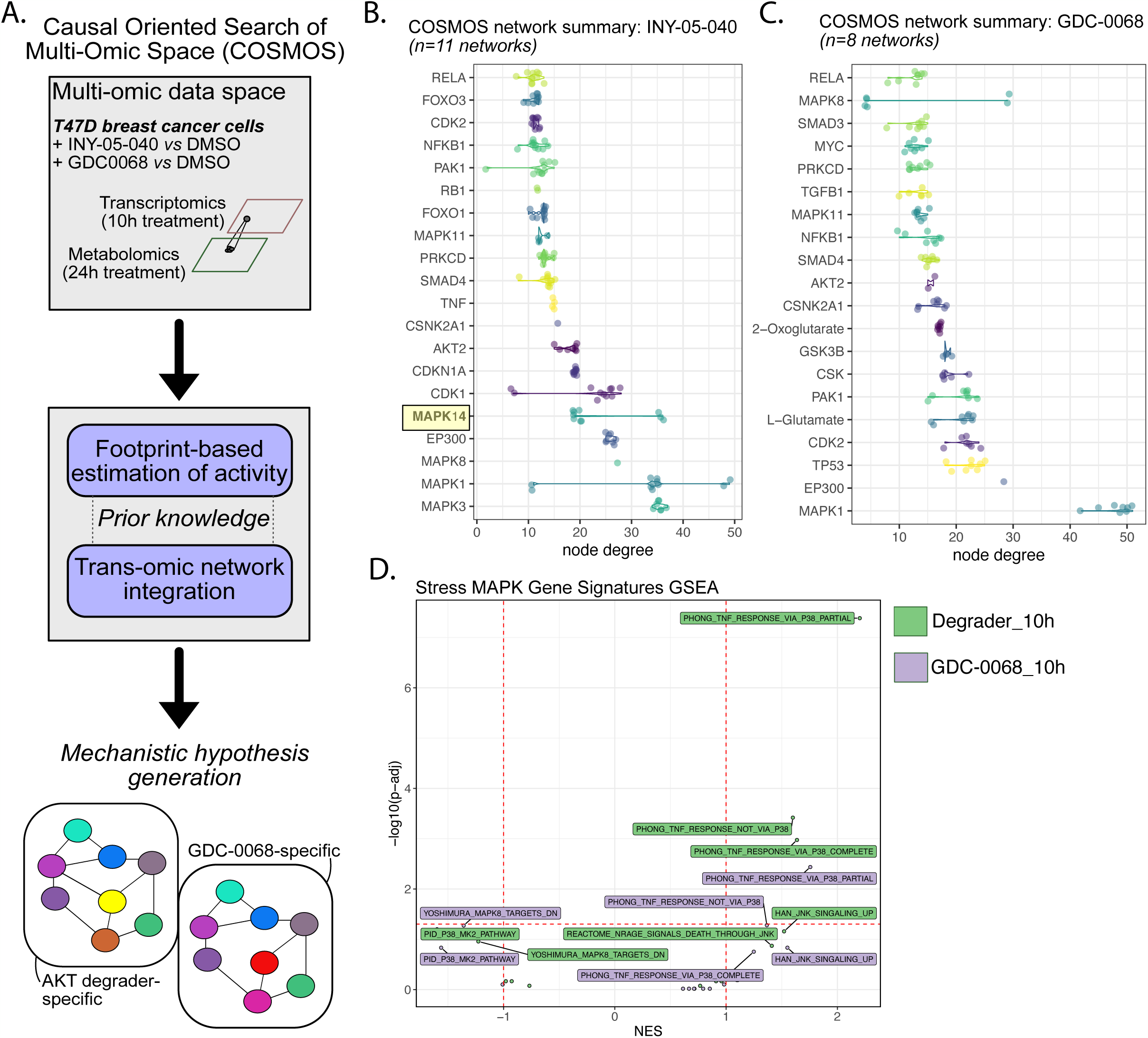
Causal Oriented Search of Multi-Omics Space (COSMOS)-based integration of transcriptomic and metabolomic datasets to identify treatment-specific networks. (**A**) Schematic illustrating the principle of COSMOS and the datasets used for multi-omic integration and predictions of treatment-specific signaling networks. (**B**) + (**C**) Top degree nodes from Degrader- and GDC-0068-specific networks plotted in increasing order. MAPK14 (protein: p38Cl) is highlighted as a Degrader network-specific top degree node. The raw COSMOS networks are included in Fig. S5 (n = 11 independent runs using Degrader data; n = 8 independent runs for GDC-0068 data). (**D**) Complementary GSEA analyses using stress MAPK- related gene sets (mSigDb C2 collection), based on the ranked *t* values from all genes for the indicated treatments relative to the corresponding DMSO treatment. Gene sets are labelled if the absolute normalized enrichment score (NES) exceeds 1.

To corroborate these findings, we next retrieved all MSigDb curated gene sets (C2 collection) featuring transcriptional changes downstream of JNK/p38 perturbation and performed GSEA using the RNAseq dataset. Three gene signatures related to TNFα signaling were positively and significantly enriched in INY-05-040-treated T47D cells after 10 h, with two of the signatures representing transcriptional changes that are either completely or partially dependent on p38 (**Fig. 3D**). These signatures originated from a study examining the response of lung cancer cells to TNFα in the presence or absence of the pan-p38 inhibitor LY479754^35^. Only one of the two p38-dependent signatures were significantly enriched for with a positive score in GDC-0068-treated cells; however, neither the significance nor the magnitude of enrichment were as strong as that observed in INY-05-040-treated cells (**Fig. 3D**). This is also consistent with a much weaker enrichment of the hallmark gene signature “TNFA_signaling_via_NFκB” in response to 10-h treatment with GDC-0068 compared to INY- 05-40 (**Fig. 2C, 2D**). Taken together, these integrated analyses point towards potent AKT degradation-induced activation of stress MAPK signaling and inflammatory gene signatures.

### Activation of stress MAPK signaling in response to AKT degradation

To validate the COSMOS predictions, we first determined the kinetics of p38/JNK signaling over a time course in a panel of breast cancer cell lines (**Fig. 4A, 4B, S6A, S6B)**. We observed potent AKT degrader-specific induction of p38α phosphorylation (T180/Y182) and the JNK target p-cJun (S73), as well as the expected increase in total c-Jun protein levels^36^ (**Fig. 4A**). Cells exhibited distinct p38/JNK signaling kinetics and magnitude in response to INY-05- 040 compared to GDC-0068. Consistently, AKT degradation resulted in more robust stress MAPK signaling induction, supporting COSMOS-based predictions.

**Figure 4.**
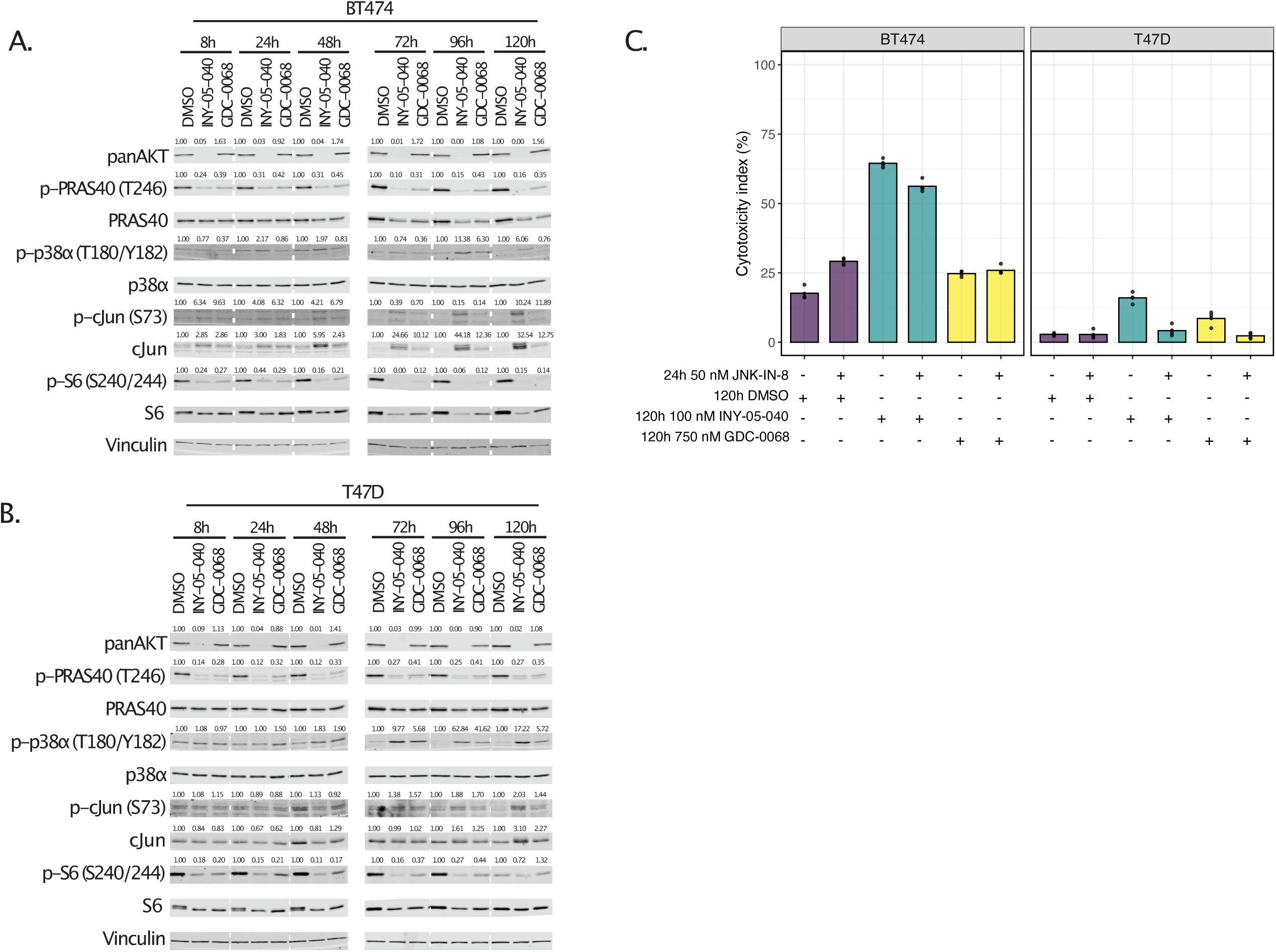
Validation of COSMOS-predicted MAPK Stress Kinase Signaling. Immunoblots for panAKT, phospho-PRAS40 (T246), total PRAS40, phospho-p38α (T180/Y182), total p38α, phospho-cJun (S73), total cJun, phospho-S6 (S240/244), total S6, and Vinculin after treatment of (**A**) BT-474 or (**B**) T47D cells for the indicated times with DMSO, 100 nM INY-05-040, or 750 nM GDC-0068. Quantification of AKT and cJun represents protein abundance over Vinculin, relative to the corresponding DMSO condition for each time point. Quantification of remaining phosphorylated proteins represent normalization to the corresponding total protein, relative to the DMSO signal for each time point. (**C**) Cytotoxicity index assayed using CellTox Green, in BT-474 or T47D cells treated for 24 h with either DMSO or 50 nM JNK-IN-8, followed by 120-h co-treatment with either DMSO, INY-05-040 (100 nM) or GDC-0068 (750 nM). The cytotoxicity index represents cytotoxicity values corrected for background fluorescence and normalized to total signal following chemical permeabilization (used as proxy measure for total cell number). For conditions of interest, the following statistics were generated using bootstrap-coupled estimation: unpaired mean percentage-point difference of JNK-IN-8 versus DMSO = 11.5 [95% CI: 8.89; 13.2]; unpaired mean percentage-point difference of INY-05-040 + JNK-IN-8 versus INY-05-040 = -8.31 [95% CI: -10.1; -5.75]. Additional supporting data related to this figure are included in Figs. S6, S7.

Among the different breast cancer cell lines examined, we found that the ER-positive cell line, BT-474, exhibited a near-binary difference in stress MAPK activation in response to AKT degradation compared to catalytic inhibition. Following 48-h treatment with INY-05-040 but not GDC-0068, BT-474 cells exhibited strong induction of p-cJun (S73) and cJun, which was sustained for at least 72 h (**Fig. 4A**). We hypothesized that induction of stress MAPK signaling contributes to AKT degrader-associated cytotoxicity. To test this, BT-474 and T47D cells were pre-treated with a low-dose (50 nM) of the covalent JNK1/2/3 inhibitor JNK-IN-8 for 24 h, followed by the addition of either GDC-0068 or INY-05-040 for another 120 h. The two cell lines were chosen as models for a potent (BT-474) versus low (T47D) cytotoxic response to AKT degradation, while both had a substantially lower magnitude of GDC-0068-induced cell death.

Consistently, the INY-05-040-induced cytotoxic response in T47D cells was fully neutralized by JNK inhibition (**Fig. 4C**). In BT-474, however, combined AKT degradation and JNK inhibition only led to a small, partial rescue of cytotoxicity (**Fig. 4C**); the increased levels of cleaved-PARP, a marker of apoptosis, in BT-474 cells treated with AKT degrader were also not reduced by co-treatment with JNK-IN-8 (**Fig. S7B**). We therefore conclude that acute and sustained stress MAPK activation is a marker of potent suppression of AKT signaling downstream of INY-05-040, but that this mechanism is not sufficient to explain the cytotoxic effect of AKT degradation.

### Global cell line screening identifies stress MAPK-associated resistance biomarkers

Given the improved cellular potency of INY-05-040, including robust downstream transcriptional and metabolic changes, we next undertook global cancer cell line profiling to determine whether INY-05-040 causes more potent growth suppression relative to GDC-0068 and the first-generation AKT degrader INY-03-041. Across 288 cancer cell lines, spanning a total of 18 different cancer lineages, INY-05-040 exhibited superior growth-inhibitory activity (**Fig. S8A**). This was based on calculation of the drug concentration required to reduce overall growth by 50 % (GI50adj, **Fig. 5A**), which includes adjustment for cell number at the start of the assay^37^. Notably, while GI50adj calculation was possible for all cell lines treated with the second-generation degrader, and for 282 cell lines treated with the first-generation degrader, this was not possible for 161 cell lines treated with GDC-0068 due to lack of sufficient growth suppression (**Fig. S8A**). Consequently, the median GI50adj value for GDC-0068 in our screen is higher than 10 µM, compared to 1.1 µM for INY-05-040 and 3.1 µM for INY-03-041.

**Figure 5.**
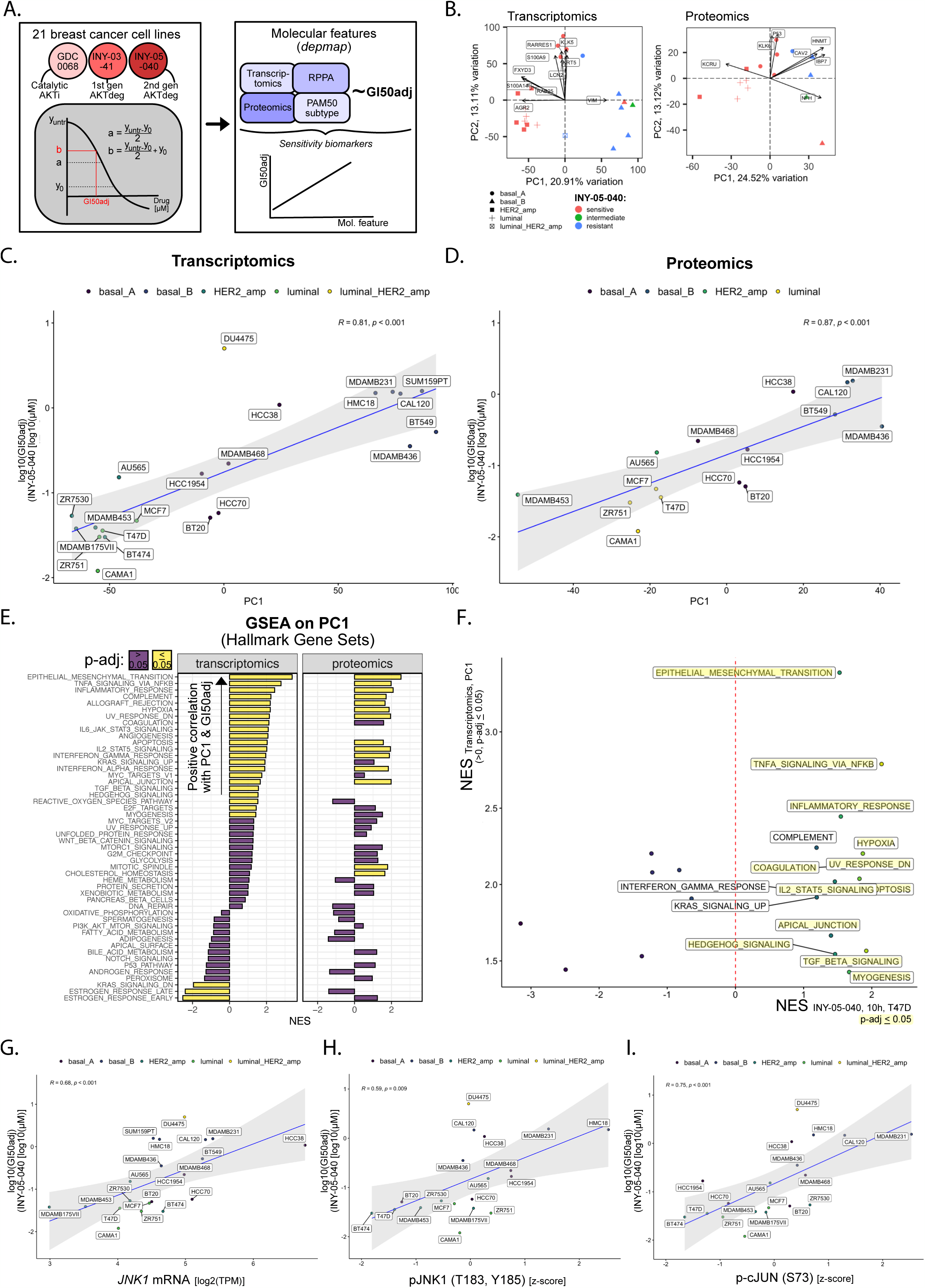
Integration of cell line screen data with publicly available omics datasets to identify sensitivity biomarkers for INY-05-040. (**A**) Analytical workflow for cell line screen processing and subsequent integration of the growth response metric (GI50adj) with publicly available cell line omics data from the DepMap project. A total of 288 cancer cell lines were profiled with GDC-0068, INY-03-41 and INY-05-040, with the full set of responses included in Fig. S8A. Subsequent integrative analyses focused on breast cancer cell lines only. Note that the applied growth response metric (GI50adj) takes into account cell line growth which is a known confounder in drug sensitivity measurements^67^. The final output corresponds to the concentration of drug that results in 50 % cell growth inhibition. (**B**) PCA on breast cancer- specific transcriptomics and proteomics data, with coloring according to sensitivity to INY-05- 040 (sensitive: GI50adj < 0.5 µM; intermediate: 0.5 µM ≤ GI50adj ≤ 1 µM; resistant: GI50adj > 1 µM; see also Fig. S8B). The PAM50 subtype of each cell line is specified by shape. Transcripts and proteins contributing the most to the observed data structure alongside PC1 and PC2 are labelled. (**C**) + (**D**) Spearman’s correlation analysis of PC1 values for each cell line and the corresponding GI50adj value for INY-05-040. A linear regression line with 95% confidence intervals (shaded area) is included in each analysis, demonstrating that cell line-specific PC1 scores can be used as proxy measures for INY-04-050 sensitivity (i.e., the higher the PC1 score, the more resistant the cell line). (**E**) GSEA (mSigDb HALLMARK gene sets) using transcript and protein loading values alongside the respective PC1, a proxy measure for sensitivity to INY-04-050; FDR ≤ 0.05. NES: normalized enrichment score. (**F**) A plot of all gene sets that were significantly and positively enriched across PC1 loadings from the DepMap transcriptomic data, and the corresponding NES from the T47D dataset following 10 h treatment with INY-05-040 (see also Fig. 2). Highlighted gene signatures were also statistically significant (FDR ≤ 0.05) in the T47D dataset. (**G**) + (**H**) + (**I**) Spearman’s correlation analysis of *JNK1* mRNA expression (G), pJNK1 (T183/Y187) (H) and p-cJun (S73) with the cell line-specific GI50adj value for INY-05-040. A linear regression line with 95% confidence intervals (shaded area) is included in each analysis. Reverse phase protein phosphorylation (RPPA) data were obtained from the DepMap project and subset for the signals of interest. Additional supporting data related to this figure are included in Fig. S8.

To identify functional biomarkers predictive of sensitivity to INY-05-040 in the 21 breast cancer cell lines profiled, we took advantage of the measured GI50adj values and the corresponding baseline transcriptomic, proteomic and reverse phase protein array (RPPA) data publicly available through the Cancer Dependency Map project (**Fig. 5A**)^38, 39^. We classified breast cancer cells into sensitive, intermediate, and resistant if the measured GI50adj was less than 0.5 µM, between 0.5 to 1 µM, and higher than 1 µM (**Fig. S8B**), respectively. Subsequent unsupervised PCA using either transcriptomic or proteomic datasets revealed a notable separation of INY-04-050-sensitive *versus* -resistant breast cancer cells, which was not simply driven by ER expression as assessed by PAM50 status (**Fig. 5B**). Except for one mixed-subtype HER2-amplified-luminal breast cancer cell line (DU4475), all examined HER2-amplified and luminal breast cancer cells were sensitive to INY-05-040. This was also observed for 4 out of 5 breast cancer cells belonging to the basal A subtype. By contrast, only 1 out of 6 basal B breast cancer cell lines were sensitive to INY-05-040, with 4 out of 6 exhibiting overt resistance (**Fig. 5B**).

Using the PC1 loadings from the transcriptomic and proteomic data, we then correlated these to the measured GI50adj values. This revealed strong and statistically significant correlations for either comparison, with higher PC1 loadings associated with higher GI50adj values and thus resistance to INY-05-040 (**Fig. 5C, 5D**). To identify the underlying molecular features, we performed GSEA on the two PC1 loadings (transcriptomic and proteomic data).

Gene sets that were positively enriched for alongside either PC1 were highly concordant and characterized by strong enrichment for epithelial mesenchymal transition and inflammatory signaling (**Fig. 5E**). Strikingly, most of these positive enrichments overlapped with those observed upon acute 10-h treatment of T47D breast cancer cells with INY-05-040 (**Fig. 5F**). Based on our mechanistic data on acute JNK activation and sensitivity to INY-05-040, we reasoned that the correlation between inflammatory gene signatures and INY-05-040 resistance in the breast cancer cell panel may reflect an already high baseline JNK activation and thus stress MAPK signaling. Accordingly, we found that both *JNK1* mRNA levels (**Fig. 5G**), JNK1 phosphorylation (T183/187) (**Fig. 5H**) and cJun phosphorylation (S73) (**Fig. 5I**) exhibited a positive and statistically significant correlation with INY-05-040 GI50adj values. Importantly, the BT-474 breast cancer cell line, which exhibits a cytotoxic response to INY-05-040 (**Fig. 4C**), had the lowest GI50adj value and the lowest values for markers of baseline JNK1 activation.

Taken together, these data demonstrate superior potency of INY-05-040-induced AKT degradation over catalytic inhibition across cancer cell lines, with low baseline levels of stress MAPK signaling correlating with heightened AKT degrader sensitivity in breast cancer cells.

## Discussion

Targeted protein degradation has emerged as both a novel therapeutic approach and a powerful experimental tool to evaluate the effects of acute protein depletion on cellular networks. Here we have reported the development of a potent and highly selective second-generation pan-AKT degrader, INY-05-040, which we used as a tool to uncover novel AKT biology. Using a multi-omic approach in breast cancer cell models, we found that AKT degradation led to unique transcriptomic and metabolomic changes, strong suppression of downstream AKT signaling, concomitant with potent activation of stress MAPK signaling. Furthermore, low baseline levels of JNK activation were associated with increased sensitivity to AKT degradation across a panel of breast cancer cell lines.

The ongoing search for targeted agents to treat patients with PI3K pathway hyperactivation has focused on the identification of more selective compounds, effective combinations to limit toxicity and improvements in patient selection^40^. Recent efforts have led to the development of the PI3Kα-selective inhibitor alpelisib (PIQRAY®), approved for the treatment of advanced hormone receptor-positive, HER2-negative breast cancer, in combination with the ER antagonist fulvestrant^41^. Alpelisib (VIJOICE®) was also recently approved for the treatment of developmental overgrowth disorders collectively known as *PIK3CA*-related overgrowth spectrum (PROS)^42, 43^. Despite this progress, both cancers and diseases of PI3K pathway activation urgently need an expansion of available treatment options to address issues of resistance and/or poor tolerability. Independent lines of evidence, including the current study, indicate that targeted protein degradation of PI3K pathway components may represent a novel therapeutic strategy, with the added benefit of sustained inhibition of downstream signaling^18, 44, 45^. This property may partly be explained by the inability of various negative feedback mechanisms within the PI3K/AKT pathway to overcome inhibition when a critical downstream transducer is absent. Prolonged cellular stress can also suppress AKT/mTORC1 activity, alongside a more complete shutdown of protein translation, which may contribute to a self-sustained feedforward loop of continued suppression of AKT signaling despite removal of the AKT degrader. This hypothesis is supported by washout experiments in which pathway reactivation was not observed or remained low for at least 72 h after degrader removal, in stark contrast to the corresponding findings with catalytic AKT inhibition with GDC-0068.

Using a network biology framework, COSMOS, we demonstrate how systematic integration of a prior knowledge with context-specific transcriptomic and metabolomic data can be used to identify and subsequently test mechanistic hypotheses on AKT degradation-selective signaling outcomes. This approach identified the stress MAPKs, p38α (*MAPK14*) and JNK1 (*MAPK8*), as differentially activated in breast cancer cells treated with the AKT degrader INY-05-040. The observed quantitative differences would have been challenging to resolve with conventional approaches, emphasizing the power of computational integration of multi-omics data and temporal analyses.

We posit that both the duration and intensity of AKT pathway inhibition is critical for eliciting the potent stress MAPK response observed with INY-05-040. At present, the precise mechanistic link between AKT degradation and stress MAPK activation remains undescribed. We speculate that ribosomal stress may contribute to the induction of stress MAPKs, since AKT and mTORC1 promote ribosome biogenesis through transcriptional and translational mechanisms. Conversely, disruption of any given step in ribosome biogenesis has been shown to cause ribosomal stress^46^. Accordingly, AKT degradation but not catalytic inhibition led to a potent and sustained reduction in total ribosomal S6 protein, which would be consistent with the low stability of ribosomal proteins in the absence of functional ribosome formation^46, 47^.

Aberrant cholesterol metabolism may also contribute to the cellular stress observed upon AKT degradation. Low cholesterol has previously been linked to increased NFκB activation and cell death in fibroblasts through a p38 MAPK-dependent mechanism^48, 49^; accordingly, activating transcriptomic signatures for both inflammatory and stress MAPK pathways were strongly enriched for in AKT degrader-treated cells. Additional studies are required to understand this putative crosstalk.

The involvement of stress MAPK and inflammatory signaling in the cellular response to AKT degradation was further supported by integration of growth inhibition measurements with publicly available omics data. The observation that the same transcriptional and signaling signatures induced upon degrader treatment of T47D cells are already elevated at baseline in breast cancer cell lines with lower sensitivity to INY-04-050 suggests that low baseline stress MAPK and inflammatory signaling activity may be a pre-requisite for potent cell growth suppression upon AKT degradation.

Several other AKT degraders have been developed to date, including the VHL-recruiting AZD5363-based AKT degrader MS21^18^. Like INY-05-040, MS21 also outperformed its parental AKT kinase inhibitor in cancer cell growth and signaling assays^18^. Additional side-by-side comparisons of MS21 and INY-05-040 are needed to determine whether both compounds share similar cellular mechanisms of action downstream of AKT degradation, given the subtle but important differences in their biochemical profiles. Interestingly, the six cell lines identified as more sensitive to AKT degradation with MS21 compared to inhibition with AZD5363 all had lower than average levels of pJNK as measured by RPPA^18^, consistent with our results.

In summary, we demonstrate vastly improved suppression of cancer cell growth with a potent second-generation AKT degrader and illustrate how protein degraders, in combination with integrated systems-level analyses, can be used to uncover novel biology that is inaccessible to conventional kinase inhibition.

## MATERIALS AND METHODS

### Synthetic Scheme of INY-05-040

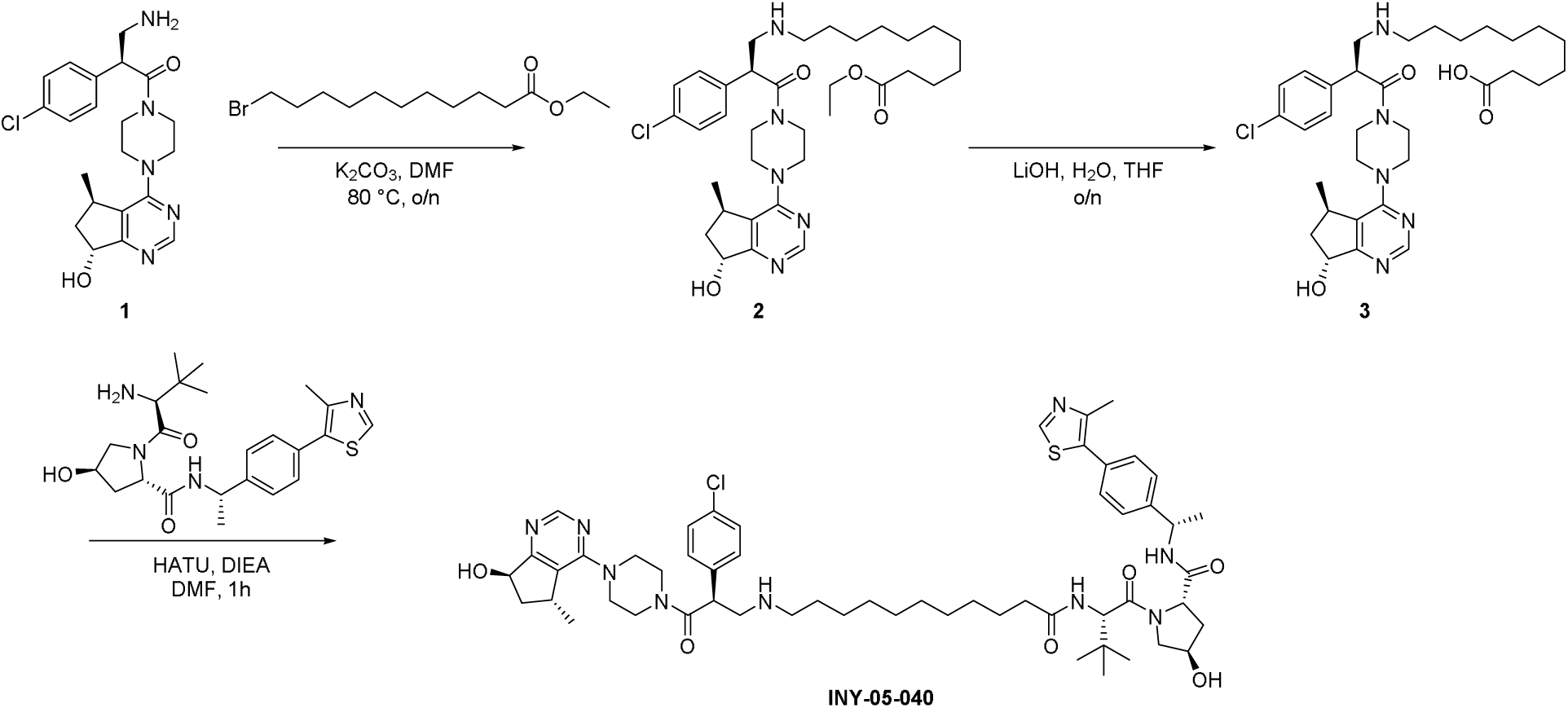

### Compound synthesis

Reagents and solvents were purchased from commercial suppliers and were used without further purification unless otherwise noted. Reactions were monitored using a Waters Acquity UPLC/MS system (Waters PDA eλ Detector, QDa Detector, Sample manager – FL, Binary Solvent Manager) using Acquity HPLC ® BEH C18 column (2.1 x 50 mm, 1.7 µm particle size): solvent gradient = 85% A at 0 min, 1% A at 1.7 min; solvent A = 0.1 % formic acid in water; solvent B = 0.1% formic acid in acetonitrile; flow rate: 0.6 mL/min. Products were purified by preparative HPLC using Waters SunFireTM Prep C18 column (19 x 100 mm, 5 µm particle size) using the indicated gradient in which solvent A = 0.05% trifluoroacetic acid (TFA) in water and solvent B = 0.05% TFA in methanol over 48 min (60 min run time) at a flow of 40 mL/min.

^1^H NMR spectra were recorded on 500 MHz Bruker Avance III spectrometer and chemical shifts are reported in million (ppm, δ) downfield from tetramethylsilane (TMS). Coupling constants (J) are reported in Hz. Spin multiplicities are described as s (singlet), br (broad singlet), d (doublet), t (triplet), q (quartet) and m (multiplet). Purities of assayed compounds were in all cases greater than 95%, as determined by reverse-phase high-performance liquid chromatography (HPLC) analysis.

#### Synthesis of INY-05-040 and INY-05-040-Neg

##### Ethyl 11-(((S)-2-(4-chlorophenyl)-3-(4-((5R,7R)-7-hydroxy-5-methyl-6,7-dihydro-5H-cyclopenta[d]pyrimidin-4-yl)piperazin-1-yl)-3-oxopropyl)amino)undecanoate (2)

(S)-3-amino-2-(4-chlorophenyl)-1-(4-((5R,7R)-7-hydroxy-5-methyl-6,7-dihydro-5H- cyclopenta[d]pyrimidin-4-yl)piperazin-1-yl)propan-1-one (150 mg, 0.36 mmol) was dissolved in DMF (2 mL). Potassium carbonate (150 mg, 1.08 mmol) was added to the reaction mixture, followed by dropwise addition of ethyl 11-bromoundecanoate (96 mg, 0.32 mmol). The reaction was stirred at 80 °C overnight. The next day, the reaction mixture was filtered and purified by reverse phase high-performance liquid chromatography (HPLC; 75% to 15% water in methanol) to obtain title compound as a yellow oil (133 mg, 56% yield). LC-MS: m/z 628.4 [M+1].

##### 11-(((S)-2-(4-chlorophenyl)-3-(4-((5R,7R)-7-hydroxy-5-methyl-6,7-dihydro-5H-cyclopenta[d]pyrimidin-4-yl)piperazin-1-yl)-3-oxopropyl)amino)undecanoic acid (3)

6 N LiOH 1 mL) and THF (tetrahydrofuran; 1 mL) were added to ethyl 11-(((S)-2-(4- chlorophenyl)-3-(4-((5R,7R)-7-hydroxy-5-methyl-6,7-dihydro-5H-cyclopenta[d]pyrimidin-4- yl)piperazin-1-yl)-3-oxopropyl)amino)undecanoate (133 mg, 0.18 mmol). The reaction mixture was stirred overnight. The next day, 1 N HCl was added to pH ∼3, and the solid was filtered and collected to obtain the title compound (128 mg, 99% yield) as a crude, which was used without further purification. LC-MS: m/z 600.42 [M+1].

##### (2S,4R)-1-((S)-2-(11-(((S)-2-(4-chlorophenyl)-3-(4-((5R,7R)-7-hydroxy-5-methyl-6,7-dihydro-5H-cyclopenta[d]pyrimidin-4-yl)piperazin-1-yl)-3-oxopropyl)amino)undecanamido)-3,3-dimethylbutanoyl)-4-hydroxy-N-((S)-1-(4-(4-methylthiazol-5-yl)phenyl)ethyl)pyrrolidine-2-carboxamide (INY-05-040)

(2S,4R)-1-((S)-2-amino-3,3-dimethylbutanoyl)-4-hydroxy-N-((S)-1-(4-(4-methylthiazol-5- yl)phenyl)ethyl)pyrrolidine-2-carboxamide (81 mg, 0.17 mmol), HATU (hexafluorophosphate azabenzotriazole tetramethyl uronium; 64 mg, 0.17 mmol), DIEA (N,N-diisopropylethylamine; 200 µL, 1.18 mmol), and DMF (dimethylformamide; 1 mL) were added to 11-(((S)-2-(4- chlorophenyl)-3-(4-((5R,7R)-7-hydroxy-5-methyl-6,7-dihydro-5H-cyclopenta[d]pyrimidin-4- yl)piperazin-1-yl)-3-oxopropyl)amino)undecanoic acid (120 mg, 0.17 mmol). The reaction was stirred for 1 h, after which the reaction was purified by reverse-phase HPLC (80% to 20% water in methanol) to obtain INY-05-040 (40 mg, 22% yield). ^1^H NMR (500 MHz, DMSO) δ 9.27 (s, 1H), 9.06 (s, 1H), 8.88 (s, 1H), 8.77 (s, 1H), 8.40 (d, J = 7.8 Hz, 1H), 7.78 (d, J = 9.3 Hz, 1H), 7.49 – 7.43 (m, 3H), 7.41 – 7.38 (m, 3H), 5.30 (t, J = 7.9 Hz, 1H), 4.92 (p, J = 7.2 Hz, 1H), 4.75 (dd, J = 8.8, 4.8 Hz, 1H), 4.52 (d, J = 9.3 Hz, 1H), 4.43 (t, J = 8.0 Hz, 1H), 4.31 – 4.25 (m, 1H), 4.03 (d, J = 42.1 Hz, 2H), 3.91 – 3.78 (m, 3H), 3.72 – 3.50 (m, 6H), 3.45 – 3.33 (m, 1H), 3.08 – 3.02 (m, 1H), 2.90 – 2.82 (m, 2H), 2.47 (s, 3H), 2.29 – 2.20 (m, 1H), 2.19 – 1.99 (m, 4H), 1.83 – 1.76 (m, 1H), 1.67 – 1.59 (m, 2H), 1.55 – 1.40 (m, 3H), 1.38 (d, J = 7.0 Hz, 2H), 1.24 (s, 12H), 1.11 (dd, J = 14.1, 6.9 Hz, 3H), 0.94 (s, 9H). LC-MS: m/z 1026.6 [M+1].

##### (2R,4S)-1-((S)-2-(11-(((S)-2-(4-chlorophenyl)-3-(4-((5R,7R)-7-hydroxy-5-methyl-6,7-dihydro-5H-cyclopenta[d]pyrimidin-4-yl)piperazin-1-yl)-3-oxopropyl)amino)undecanamido)-3,3-dimethylbutanoyl)-4-hydroxy-N-((S)-1-(4-(4-methylthiazol-5-yl)phenyl)ethyl)pyrrolidine-2-carboxamide (INY-05-040-Neg)

INY-05-040-Neg was synthesized using similar procedures as INY-05-040 using (2R,4S)-1-((S)-2-(11-(((S)-2-(4-chlorophenyl)-3-(4-((5R,7R)-7-hydroxy-5-methyl-6,7-dihydro-5H- cyclopenta[d]pyrimidin-4-yl)piperazin-1-yl)-3-oxopropyl)amino)undecanamido)-3,3- dimethylbutanoyl)-4-hydroxy-N-((S)-1-(4-(4-methylthiazol-5-yl)phenyl)ethyl)pyrrolidine-2- carboxamide as the starting material. ^1^H NMR (500 MHz, DMSO) δ 8.99 (s, 1H), 8.70 (s, 1H), 8.54 (s, 1H), 8.40 (s, 1H), 8.06 (d, J = 8.0 Hz, 1H), 7.89 (d, J = 7.8 Hz, 1H), 7.49 (d, 2H), 7.45 – 7.44 (m, 3H), 7.38 – 7.35 (m, 2H), 5.20 (t, J = 7.6 Hz, 1H), 4.91 (p, 1H), 4.48 (dd, J = 8.7, 5.0 Hz, 1H), 4.42 – 4.37 (m, 2H), 4.33 – 4.28 (m, 1H), 3.98 (s, 1H), 3.80 (dd, J = 10.4, 5.4 Hz, 2H), 3.77 – 3.58 (m, 6H), 3.56 – 3.45 (m, 3H), 3.14 – 3.07 (m, 1H), 2.94 – 2.86 (m, 2H), 2.47 (s, 3H), 2.29 – 2.21 (m, 1H), 2.14 – 1.93 (m, 6H), 1.62 – 1.52 (m, 2H), 1.52 – 1.38 (m, 3H), 1.32 (d, J = 7.0 Hz, 3H), 1.26 – 1.17 (m, 13H), 1.08 (d, J = 6.9 Hz, 3H), 0.97 (s, 9H). LC-MS: m/z 1026.57 [M+1].

### Biochemical Selectivity Assay

Biochemical selectivity across 468 kinases was measured through the scanMAX kinase assay panel provided through Eurofins Discovery.

### Cell Culture

BT-474, T47D, MCF-7, and MDA-MB-468 cells were obtained from ATCC and cultured in RPMI media supplemented with 10% heat inactivated fetal bovine serum without antibiotics at 37 °C in the presence of 5 % CO_2_. Cells were maintained in Corning TC-treated 15 cm culture dishes (Corning Cat. # 08-772-24) in 20 mL medium. Medium was replenished every 3 days, until cells reached 70-90% confluence. To passage, cells were washed once with 10 mL PBS and incubated for 5-10 min at 37 °C with 0.25% Trypsin 0.1% EDTA and passaged up to 5 times in the same dish. Cells were maintained in culture for up to one month. Cells were routinely tested for mycoplasma using a Mycoplasma Detection Kit (Lonza Cat. # LT07-218).

### Drug Treatment Experiments

#### Cell Death Rescue Experiments

T47D and BT-474 cells were plated at 4,000 or 6,000 cells/well in 80 μl RPMI + 10% FBS medium in black-walled clear-bottom 96-well plates (Fisher Cat # 12-566-70), medium was exchanged the following day with 90 μl medium plus 10 μl of drug containing medium for a 24-h pre-treatment. The following day, medium was exchanged with 80 μl complete medium plus 20 μl of drug containing medium. After an additional three days, 100 μl of drug-containing medium was replenished without removing existing medium, to prevent nutrient depletion until assay endpoint.

#### Biochemical Signaling

Depending on the length of the experiment, cells were plated at 150,000-250,000 cells/mL (MDA-MB-468), 200,000-300,000 cells/mL (BT-474, T47D) in RPMI medium with 10% serum at 2 mL per well in 6-well treated tissue culture plates (Greiner, Cat. # TCG-657160) and incubated overnight. The next day, medium was exchanged, and cells were treated with the indicated compounds at the appropriate concentration and protein lysates were harvested at the times specified. Time courses were conducted in reverse by drugging the longest time point first, followed by subsequent time points such that all samples were collected at the same time. At the time of harvest, cells were washed once with 2 mL of ice-cold PBS and either snap frozen on dry ice and stored at -80 °C, or harvested immediately (see “Immunoblotting”).

#### RNAseq

Cells were plated at 300,000 cells/mL in RPMI medium with 10% serum at 2 mL per well in 6-well treated tissue culture plates (Greiner, Cat. # TCG-657160) to achieve 75% density the following day and incubated overnight. The following day, media stocks containing indicated compounds were prepared and used to treat all conditions at respective time points, stored at 4°C between treatments. After 5 h and 10 h of treatment, wells were washed once with 2 mL ice cold PBS and aspirated completely, snap frozen on dry ice, and stored at -80 °C until all replicates were collected. Three independent biological replicates were plated ion sequential days. See “RNA sequencing” for RNA harvest protocol. In parallel, samples were also collected for protein harvest for confirmation of consistent drug effect on cellular signaling.

#### PRO-seq

T47D cells were seeded at 8 x 10^6^ cells per 15 cm plates (Corning Cat. # 08-772-24) in 15 mL medium for PRO-seq samples, or at 3 x 10^6^ cells per 10 cm plates (Westnet Cat. # 353003) in 8 mL medium for protein samples to achieve 70% confluence. The following day a stock of drug-containing medium was prepared and used to treat technical triplicates for each condition. Technical triplicate protein replicates treated the same way were collected in parallel. Protein plates were washed once with 8 mL of ice-cold PBS and snap frozen on dry ice, then stored at -80 °C until all replicates were collected (see “Immunoblotting”).

#### Metabolomics

Cells were plated at 2 x 10^6^ cells/plate in RPMI medium with 10% in 3 mL per plate in 60 mm treated tissue culture plates (Corning, Cat. # 430166) and incubated overnight. The next day, stocks of medium were prepared containing the indicated compounds at the appropriate concentration, and medium was exchanged for drug-containing medium. Three independent biological replicates were performed for metabolomics experiments, each comprising technical triplicates for metabolite plates and technical duplicate of parallel protein samples used to assess suppression of signaling and normalize metabolite levels to total protein content. Due to an apparent loss of potency, the dose of GDC-0068 was increased to 750 nM in Trial 3, compared to 500 nM in Trials 1 and 2, to ensure consistent biochemical signaling suppression across all runs.

### Proliferation Assays

T47D, MDA-MB-468, MCF-7 or BT-474 cells were plated in 384 well plates at 250 cells per well. After 24 h, cells were treated with GDC-0068, AZD5363, MK-2206, ARQ-092, INY-03- 041, INY-05-040, INY-05-040-Neg, or VH032 compounds at concentrations indicated for 72 h. The anti-proliferative effects of these compounds were assessed using the Cell Titer Glo assay kit (Promega Cat. # G7570) following manufacturer protocol. EC_50_ values were determined using GraphPad Prism using nonlinear regression curve fitting.

### CellTox Green Cell Death Assay

Cell viability was assayed with a CellTox Green cell death assay. Cells in 96-well plates (ThermoFisher Cat. # 165305) were treated with a 1:1000 dilution (in assay buffer) of CellTox Green dye for 30 min at room temperature, protected from light. Fluorescence intensity, corresponding to binding of CellTox Green dye to double-stranded DNA from dead cells, was measured on a SpectraMax iD3 Microplate Reader (485 nm excitation / 520 nm emission) from the bottom, with an integration time of 400 ms and 9 multi-point readings per well. To estimate the total number of cells for subsequent normalization, all wells were subsequently permeabilized with 0.1% Triton X-100 (Fisher Scientific Cat. # BP151-100), and enough CellTox Green reagent to maintain “1X” final concentration. After incubating for 30 min at room temperature, protected from light, the final fluorescence intensity was measured as above.

Readings from each well were averaged and corrected by subtracting the average background signal from wells with medium and CellTox Green and no cells. The cytotoxicity index was calculated for treatments of interest by dividing background-corrected non-permeabilized readings by the corresponding permeabilized readings to assess the percentage cell death. Each assay run was quality checked by inclusion of a standard curve of increasing cell number, followed by permeabilization and measurement of the CellTox Green signal. All raw data and annotated analysis scripts are available on the associated OSF project website (https://osf.io/fasqp/).

In parallel, cell health and CellTox Green uptake were also assessed by light microscopy, with image capture on a Keyence BZ-X800 (brightfield and 488 nm) and an ECHO Scope (brightfield only; 10X). These images were used as internal QC and are not incorporated in the final manuscript but have been deposited on the OSF project website (https://osf.io/fasqp/) as further supporting evidence.

### Immunoblotting

Cells were washed once in 1x PBS then lysed in RIPA buffer (150 mM Tris-HCl, 150 mM NaCl, 0.5% (w/v) sodium deoxycholate, 1% (v/v) NP-40, pH 7.5) containing 0.1% (w/v) sodium dodecyl sulfate, 1 mM sodium pyrophosphate, 20 mM sodium fluoride, 50 nM calyculin, and 0.5% (v/v) protease inhibitor cocktail (Sigma-Aldrich Cat. # P8340-5ML) for 15 min. Cell extracts were precleared by centrifugation at 18,800 x g for 10 min at 4 °C. The Bio-Rad DC protein assay was used to assess protein concentration as per the manufacturer’s instructions, and sample concentration was normalized using 2x SDS sample buffer. Next, 20 μg of protein lysates and PageRuler Plus (Fisher Cat. # PI26619) prestained protein ladder were resolved on 10% acrylamide gels by SDS-polyacrylamide gel electrophoresis and electrophoretically transferred to nitrocellulose membrane (BioRad Cat. # 1620112) at 100 volts for 90 min.

Membranes were blocked in 5% (w/v) nonfat dry milk (Fisher Cat# NC9022655/190915ASC) or 5% (w/v) bovine serum albumin (Boston Bioproducts Cat. # P-753) in Tris-buffered saline (TBS) for 1 h, then incubated with specific primary antibodies diluted 1:1000 in 5% (w/v) bovine serum albumin in TBS-T (TBS with 0.05% Tween-20) at 4 °C overnight, shaking. The next day, membranes were washed 3 times for 5 min each with TBS-T then incubated for 1 h at room temperature with fluorophore-conjugated secondary antibodies (LI-COR Biosciences) in 5% (w/v) nonfat dry milk, protected from light. The membrane was washed again 3 times for 5 min each with TBS-T, followed by a final 5-min wash in TBS, then imaged with a LI-COR Odyssey CLx Imaging System (LI-COR Biosciences).

For Figure S7 blots, medium containing dead or floating cells was collected from each well and centrifuged for 5 min at 300 x rcf. Medium was aspirated, and the pellet lysed in RIPA buffer and combined with protein harvested from corresponding adherent cells as described above.

Quantification was performed in ImageStudioLite Software (Licor Biosciences) by drawing rectangles around bands to capture band signal intensities: total pixel intensity minus background pixel intensity. Relative phospho-protein signal was performed for each lane by dividing phospho-protein signal intensity by corresponding total protein signal intensity, while relative AKT signal was calculated by dividing AKT signal intensity by Vinculin signal intensity. Normalization to DMSO samples was performed by dividing relative signal intensity for each condition by the corresponding DMSO signal intensity values.

### Proteomics

#### MOLT4 Cell Culture and Sample Preparation

MOLT4 cells (T lymphoblast established from a 19-year-old male patient with Acute Lymphoblastic Leukemia in relapse) were grown in RPMI-1640 media including 2mM L-glutamine (Gibco) and supplemented with 10% fetal bovine serum (Gibco) in a 37 °C incubator with 5% CO_2_. MOLT4 cells were treated with DMSO or 250 nM INY-05-040 for 4 h. Cells were harvested by centrifugation, followed by addition of lysis buffer (8 M Urea, 50 mM NaCl, 50 mM 4-(2hydroxyethyl)-1-piperazineethanesulfonic acid (EPPS) pH 8.5), 1x cOmplete protease inhibitor (Roche) and 1x PhosphoStop (Roche). Cells were subsequently homogenized by 20 passes through a 21-gauge (1.25 in. long) needle to achieve a cell lysate with a protein concentration between 0.5-4 mg/mL. The homogenized sample was clarified by centrifugation at 20,000 x g for 10 min at 4°C. A Bradford assay was used to determine the final protein concentration in the cell lysate. 200 µg protein for each sample were reduced, alkylated precipitated using methanol/chloroform and dried as previously described^50^. Precipitated protein was resuspended in 4 M Urea, 50 mM HEPES pH 7.4, followed by dilution to 1 M urea with the addition of 200 mM EPPS pH 8 for digestion with LysC (1:50; enzyme:protein) for 12 h at RT. The LysC digestion was diluted to 0.5 M Urea, 200 mM EPPS pH 8 and then digested with trypsin (1:50; enzyme:protein) for 6 h at 37°C. Tandem mass tag (TMT) reagents (Thermo Fisher Scientific) were dissolved in anhydrous acetonitrile (ACN) according to manufacturer’s instructions. Anhydrous ACN was added to each peptide sample to a final concentration of 30% v/v, and labeling was induced with the addition of TMT reagent to each sample at a ratio of 1:4 peptide:TMT label. The 11-plex labeling reactions were performed for 1.5 h at RT and the reaction quenched by the addition of 0.3% hydroxylamine for 15 minutes at RT. The sample channels were combined at a 1:1 ratio, desalted using C18 solid phase extraction cartridges (Waters) and analyzed by LC-MS for channel ratio comparison. Samples were then combined using the adjusted volumes determined in the channel ratio analysis and dried down in a speed vacuum. The combined sample was then resuspended in 1% formic acid and acidified (pH 2-3) before being subjected to desalting with C18 SPE (Sep-Pak, Waters). Samples were then offline fractionated into 96 fractions by high pH reverse-phase HPLC (Agilent LC1260) through an Aeris peptide XB-C18 column (phenomenex) with mobile phase A containing 5% acetonitrile and 10 mM NH4HCO3 in LC-MS grade H_2_O, and mobile phase B containing 90% acetonitrile and 10 mM NH4HCO3 in LC-MS grade H_2_O (both pH 8.0). The 96 resulting fractions were then pooled in a non-contiguous manner into 24 fractions and desalted using solid phase extraction plates (SOLA, Thermo Fisher Scientific) followed by subsequent mass spectrometry analysis.

#### Data Acquisition

Data were collected using an Orbitrap Fusion Lumos mass spectrometer (Thermo Fisher Scientific, San Jose, CA, USA) coupled with a Proxeon EASY-nLC 1200 LC pump (Thermo Fisher Scientific). Peptides were separated on a 50 cm and 75 μm inner diameter Easyspray column (ES803a, Thermo Fisher Scientific). Peptides were separated using a 190 min gradient of 6 – 27% acetonitrile in 1.0% formic acid with a flow rate of 300 nL/min. Each analysis used an MS3-based TMT method as described previously (McAlister et al., 2014). The data were acquired using a mass range of m/z 340 – 1350, resolution 120,000, AGC target 5 x 105, maximum injection time 100 ms, dynamic exclusion of 120 seconds for the peptide measurements in the Orbitrap. Data dependent MS2 spectra were acquired in the ion trap with a normalized collision energy (NCE) set at 35%, AGC target set to 1.8 x 104 and a maximum injection time of 120 ms. MS3 scans were acquired in the Orbitrap with a HCD collision energy set to 55%, AGC target set to 2 x 105, maximum injection time of 150 ms, resolution at 50,000 and with a maximum synchronous precursor selection (SPS) precursors set to 10.

#### Data Analysis

Proteome Discoverer 2.4 (Thermo Fisher) was used for .RAW file processing and controlling peptide and protein level false discovery rates, assembling proteins from peptides, and protein quantification from peptides. MS/MS spectra were searched against a Swissprot human database (February 2020) with both the forward and reverse sequences. Database search criteria are as follows: tryptic with two missed cleavages, a precursor mass tolerance of 10 ppm, fragment ion mass tolerance of 0.6 Da, static alkylation of cysteine (57.02146 Da), static TMT labeling of lysine residues and N-termini of peptides (229.16293 Da), variable phosphorylation of serine, threonine and tyrosine (79.966 Da), and variable oxidation of methionine (15.99491 Da). TMT reporter ion intensities were measured using a 0.003 Da window around the theoretical *m/z* for each reporter ion in the MS3 scan. Peptide spectral matches with poor quality MS3 spectra were excluded from quantitation (summed signal-to-noise across 11 channels < 100 and precursor isolation specificity < 0.5). Only proteins containing at least two unique peptides identified in the experiment were included in final quantitation. Reporter ion intensities were normalized and scaled using in-house scripts in the R framework^51^. Significant changes comparing the relative protein abundance of treatment samples to the DMSO control treatments were assessed by moderated t-test as implemented in the limma package within the R framework^52^.

### RNA sequencing

#### Sample Preparation

Snap-frozen cells were thawed on ice and RNA extracted with Takara’s Nucleospin RNA Plus kit (Takara Cat. # 740984.50) according to the manufacturer’s instructions. RNA integrity was assessed for quantity and purity by Nanodrop 1000. Samples were submitted to Novogene for integrity assessment (Agilent 2100 analysis), mRNA library preparation (unstranded), and paired-end (150 bp) sequencing on a NovaSeq S4 flow cell.

#### Raw read mapping, counting and differential expression

Raw read processing was performed with the Nextflow (version 20.07.1) nf-core RNAseq pipeline (version 1.4.2)^53^, with Spliced Transcripts Alignment to a Reference (STAR)^54^ for read alignment to the human genome (Homo_sapiens.GRCh38.96.gtf) and featureCounts^55^ for counting of mapped reads (multimapped reads were discarded).

All subsequent data processing was performed in R, with differential gene expression analysis following the limma-voom method^56^. Filtering of low gene expression counts was performed with the *TCGAbiolinks* package with quantile value 0.75 (chosen empirically based on the observed count distribution). Next, read count normalization was performed with the trimmed mean of M (TMM) method^57^. PCA was done using the *PCAtools* package. The mean-variance relationship was modelled with voom(), followed by linear modelling and computation of moderated t-statistics using the lmFit() and eBayes() functions in the *limma* package^56^. Experimental replicate was included as a batch effect term in the model. The associated p-values for assessment of differential gene expression were adjusted for multiple comparisons with the Benjamini-Hochberg method at false-discovery rate (FDR) = 0.05^58^. Adjustments were performed separately for each contrast of interest. Subsequent gene annotations were performed with *BioMart* within R^59^, using the associated ENSEMBL Gene IDs as key values. Intersection plots and heatmaps were generated using the *ComplexHeatmap* package^60^. Clustering was performed using the Ward.D2 method. Columns were clustered according to Euclidean distance, while rows (genes) were clustered according to Spearman’s correlation, i.e. patterns of change as opposed to maximum values.

#### Gene set enrichment analysis

The *msigdbr* package was used to retrieve the indicated gene signatures. GSEA was performed with the *fgsea* package^61^, using the list of all genes ranked according to their *t* statistic for a comparison of interest. The choice to use *the t* statistic ensures that the gene ranking considers signal magnitude (fold-change) as well as uncertainty of estimation.

Normalized enrichment values and associated p-values were calculated with the fgseaMultilevel() function, using default settings. The normalized enrichment score computed by the algorithm corresponds to the enrichment score normalized to mean enrichment of random samples, using the same gene set size.

#### Transcription factor footprint analysis

The voom-normalized counts were used to predict transcription factor activities with DoRothEA^25^, choosing regulons within confidence groups “A”, “B” and “C” (low-confidence regulons in groups “D” and “E” were therefore not considered). As per the developer’s recommendations, the “minsize” argument in the options was set to “5”, and “eset.filter” was set to “FALSE”. Exact details can be retrieved from the deposited code.

Annotated scripts for all analysis steps post-read processing are provided on the OSF project webpage (https://osf.io/3f2m5/).

### Precision nuclear run-on sequencing (PRO-seq)

#### Sample Preparation

To harvest cell pellets for PRO-seq, cells were washed once with 8 mL room temperature 1X PBS then trypsinized for 5 min. Trypsin was quenched with ice cold DMEM + 10% FBS and cells were collected in a 50 mL conical tube and placed onto ice immediately. Cells were spun at 300 x g for 4 min at 4°C, supernatant was removed, and cells were resuspended in 250 μL Buffer W (10 mM Tris-Cl, pH 8.0; 10 mM KCl; 250 mM Sucrose; 5 mM MgCl2; 1 mM EGTA; 0.5 mM DTT; 10 % (v/v) Glycerol; Protease inhibitor tablet (EDTA-free), 0.02% SUPERase-IN RNAse inhibitor) to obtain a single-cell suspension by pipetting. 10 mL of Buffer P (10 mM Tris-Cl, pH 8.0; 10 mM KCl; 250 mM Sucrose; 5 mM MgCl2; 1 mM EGTA; 0.1 % (v/v) Igepal CA-630; 0.5 mM DTT; 0.05 % (v/v) Tween-20; 10 % (v/v) Glycerol; Protease inhibitor tablet (EDTA-free), 0.02% SUPERase-IN RNAse inhibitor) was added and cells were incubated on ice for 5 min, then spun at 400 x g for 4 min at 4 °C. Supernatant was removed and Buffer W was added and pipetted gently 2-3 times to resuspend cell suspension. An additional 9 mL of Buffer W was added to each tube, and cells were spun at 400 x g for 4 min at 4 °C. An additional wash with Buffer W was completed as above and supernatant was carefully decanted so cell pellets were not disturbed. Pellets were resuspended in Buffer F (50 mM Tris-Cl, pH 8.0; 40 % (v/v) glycerol; 5 mM MgCl2; 1.1 mM EDTA; 0.5 mM DTT, and SUPERase-IN RNAse inhibitor) and transferred to a 1.5 mL tube. The 50 mL tube was rinsed once more with 250 μl of Buffer F and added to the corresponding 1.5 mL tube for a final volume of 500 μl per sample. 10μl was reserved for counting after dilution 1:10 and 1:20 in PBS, both with and without trypan blue to calculate the fraction of permeabilized cells. Cells were diluted to 1 x 10^6^ permeabilized cells per 100 μl and a total of 5 x 10^6^ cells were aliquoted in 500 μl of Buffer F and snap frozen in liquid nitrogen and stored at -80°C until further processing. RNAse-free water was used to make all reagents and solutions, and solutions were filter sterilized with 0.2 μM filters into RNAse-free plastic bottles. Two independent biological replicates were collected, alongside the corresponding protein samples to confirm drug action at the signaling level.

#### PRO-seq library construction

Aliquots of frozen (-80 °C) permeabilized cells were thawed on ice and pipetted gently to fully resuspend. Aliquots were removed and permeabilized cells were counted using a Luna II, Logos Biosystems instrument. For each sample, 1 million permeabilized cells were used for nuclear run-on, with 50,000 permeabilized *Drosophila* S2 cells added to each sample for normalization. Nuclear run-on assays and library preparation were performed essentially as described in Reimer et al.^62^ with the following modifications: 2X nuclear run-on buffer consisted of (10 mM Tris (pH 8), 10 mM MgCl2, 1 mM DTT, 300 mM KCl, 40uM/ea biotin-11-NTPs (Perkin Elmer), 0.8 U/µL SuperaseIN (Thermo), 1% sarkosyl). Run-on reactions were performed at 37 °C. Adenylated 3’ adapter was prepared using the 5’ DNA adenylation kit (NEB) and ligated using T4 RNA ligas’ 2, truncated KQ (NEB, per manufacturer’s instructions with 15% PEG-8000 final) and incubated at 16 °C overnight. 180 µL of betaine blocking buffer (1.42 g of betaine brought to 10 mL with binding buffer supplemented to 0.6 uM blocking oligo (TCCGACGATCCCACGTTCCCGTGG/3InvdT/)) was mixed with ligations and incubated 5 min at 65°C and 2 min on ice prior to addition of streptavidin beads. After T4 polynucleotide kinase (NEB) treatment, beads were washed once each with high salt, low salt, and blocking oligo wash (0.25X T4 RNA ligase buffer (NEB), 0.3 µM blocking oligo) solutions and resuspended in 5’ adapter mix (10 pmol 5’ adapter, 30 pmol blocking oligo, water). The 5’ adapter ligation was per Reimer et al.^62^ but with 15% PEG-8000 final. Eluted cDNA was amplified 5-cycles (NEBNext Ultra II Q5 master mix (NEB) with Illumina TruSeq PCR primers RP-1 and RPI-X) following the manufacturer’s suggested cycling protocol for library construction. The product (preCR) was serially diluted and used for test amplification to determine the optimal PCR conditions for the final libraries. The pooled libraries were paired-end sequenced using the Illumina NovaSeq platform.

#### PRO-seq raw read processing

All custom scripts described herein are available on the AdelmanLab Github (https://github.com/AdelmanLab/NIH_scripts). Using a custom script (trim_and_filter_PE.pl), FASTQ read pairs were trimmed to 41bp per mate, and read pairs with a minimum average base quality score of 20 retained. Read pairs were further trimmed using cutadapt 1.14 to remove adapter sequences and low-quality 3’ bases (--match-read-wildcards -m 20 -q 10). R1 reads, corresponding to RNA 3’ ends, were then aligned to the spiked in Drosophila genome index (dm3) using Bowtie 1.2.2 (-v 2 -p 6 –best –un), with those reads not mapping to the spike genome serving as input to the primary genome alignment step (using Bowtie 1.2.2 options -v 2 –best). Reads mapping to the hg38 reference genome were then sorted, via samtools 1.3.1 (-n), and subsequently converted to bedGraph format using a custom script (bowtie2stdBedGraph.pl) that counts each read once at the exact 3’ end of the nascent RNA. Because R1 in PRO-seq reveals the position of the RNA 3’ end, the “+” and “-“ strands were swapped to generate bedGraphs representing 3’ end positions at single nucleotide resolution.

Annotated transcription start sites were obtained from human (GRCh38.99) GTFs from Ensembl. After removing transcripts with {immunoglobulin, Mt_tRNA, Mt_rRNA} biotypes, PRO-seq signal in each sample was calculated in the window from the annotated TSS to +150 nt downstream, using a custom script, make_heatmap.pl. Given good agreement between replicates and similar return of spike-in reads, bedGraphs were merged within conditions, and depth-normalized, to generate bigwig files binned at 10 bp.

#### Refinement of gene annotation (GGA) using PRO-seq and RNAseq

The corresponding paired-end RNA-seq reads were mapped to the hg38 reference genome via HISAT2 v2.2.1 (--known-splicesite-infile). To select gene-level features for differential expression analysis, and for pairing with PRO-seq data, we assigned a single, dominant TSS and transcription end site (TES) to each active gene. This was accomplished using a custom script, get_gene_annotations.sh (available at https://github.com/AdelmanLab/GeneAnnotationScripts), which uses RNAseq read abundance and PRO-seq R2 reads (RNA 5’ ends) to identify dominant TSSs, and RNAseq profiles to define most commonly used TESs. RNAseq and PRO-seq data from all conditions were used for this analysis, to comprehensively capture gene activity in these samples.

Reads were summed within the TSS to TES window for each active gene using the make_heatmap script (https://github.com/AdelmanLab/NIH_scripts), which counts each read once, at the exact 3’ end location of the nascent RNA.

#### Differential expression analysis

All subsequent processing of the PRO-seq count data were as described above for the RNAseq count data. Filtering of low counts was performed with the *TCGAbiolinks* package with quantile value 0.1. Annotated scripts for the associated analysis steps are provided on the OSF project webpage (https://osf.io/3f2m5/).

### Metabolomics

#### Sample Preparation

For metabolite extraction, media was aspirated, and cells were washed once with ice- cold PBS on wet ice. Ice-cold 80% (v/v) mass spec-grade methanol was added, the plate was transferred to dry ice and scraped, and the resulting solution was collected. Protein samples were collected in duplicate for normalization to protein content and signaling validation as described above. Insoluble material was pelleted by centrifugation at 20,000 x g for 5 min, and the resulting supernatant was evaporated under nitrogen gas. Samples were resuspended in 20 ml HPLC-grade water for LC/MS analysis.

#### Data Acquisition

For polar metabolite profiling, 5 µl from each sample were injected and analyzed using a 5500 QTRAP hybrid triple quadrupole mass spectrometer (AB/SCIEX) coupled to a Prominence UFLC HPLC system (Shimadzu) with HILIC chromatography (Waters Amide XBridge), via selected reaction monitoring (SRM) with polarity switching. A total of 295 endogenous water- soluble metabolites were targeted for steady-state analyses. Electrospray source voltage was +4950 V in positive ion mode and −4500 V in negative ion mode. The dwell time was 3 ms per SRM transition 32. Peak areas from the total ion current for each metabolite were integrated using MultiQuant v2.1.1 software (AB/SCIEX).

#### Data Analysis

Prior to differential abundance analysis, the raw metabolomics data were preprocessed as follows. Untrusted metabolites were removed from the datasets; these included: SBP, shikimate, shikimate-3-phosphate, spermidine, spermine, succinyl-CoA-methylmalonyl-CoA- nega, trehalose-6-phosphate, trehalose-sucrose, malonyl-CoA-nega, N-acetyl spermidine, N- acetyl spermine, Acetylputrescine, NAD+_nega, NADH-nega, NADP+_nega, NADPH-nega, O8P-O1P, OBP, propionyl-CoA-neg, putrescine, acetoacetyl-CoA_neg, acetyl-CoA_neg,

Cellobiose, coenzyme A_nega, glutathione, glutathione disulfide-posi. Next, metabolites with low peak intensities (<10,000) across at least 50% of the samples were removed. Finally, all metabolites with 0 intensity in more than 3 samples were also removed; any metabolites with 0 intensity in < 3 samples were removed in the final differential abundance analysis steps.

Metabolomics data normalized to matched protein samples from three independent experiments, each including three separate cell cultures per treatment, were combined into one dataset. Metabolites with missing (“NA”) or negative values in at least one trial were removed, resulting in 169 metabolites included in the final analyses. These were processed for differential abundance testing using the limma-voom method (see RNAseq *Data Analysis*), with quantile normalization due to significant heteroscedascity. Subsequent linear modelling and computation of moderated *t*-statistics was performed with *lmFit()* and *eBayes()* as for the RNAseq data, including experimental replicate as blocking factor due to a noticeable batch effect. Heatmap generation and clustering of differentially abundant metabolites was performed as described for the RNAseq data. Annotated scripts for the associated analysis steps are provided on the OSF project webpage (https://osf.io/3f2m5/).

### Causal Oriented Search of Multi-Omic Space (COSMOS)

The RNAseq input data for COSMOS consisted of transcription factor *t* values from DoRothEA and the *limma-voom*-based *t* statistic for all genes, irrespective of significance, for a given contrast of interest (GDC-0068 *versus* DMSO; INY-05-040 *vs* DMSO). The latter served as additional constraints on the solver. Metabolite data for COSMOS consisted of the *limma-voom*-based *t* statistic for metabolites with unadjusted p-value ≤ 0.05, resulting in 58 metabolites for GDC-0068 and 77 metabolites for INY-05-040. The decision to use unadjusted p-values for filtering was made *a priori* due to well-known fact of high correlation across groups of metabolites, thus making the resulting corrections for multiple comparisons overly restrictive.

Metabolite names had to be mapped to their corresponding PubChem ID, which was facilitated by the R packages *KEGGREST* and *webchem*^63^.

Exact code for generation of both RNAseq and metabolite values in the correct format for COSMOS, as well as extensive details on all required installations and subsequent code for running COSMOS on a high-performance computer cluster, are provided on the accompanying OSF project page (https://osf.io/tdvur/). Briefly, the algorithm relies on CARNIVAL’s Integer Linear Programming (ILP) optimization, which was rerun multiple times for each dataset to determine the most consistent network predictions. Settings for each run, including the resulting network gap values, are provided in an accompanying table on the OSF project page.

Differences included explicit indication of AKT1/2 inhibition (AKT3 was not expressed in T47D cells) as well as shuffling of individual *t* values for the background transcriptome, thus artificially forcing the solver to initiate the optimization from different starting points.

A “forward” optimization run to connect deregulated transcription factors (“signaling” input) as starting points to metabolites was performed first, followed by a “backward” optimization run connecting metabolites to signaling components. These optimization runs were used as the basis for the actual forward and final runs defining the output of the algorithm. Time limits for solving were set empirically, ensuring that the gap values of the resulting networks were ≤ 5% (indicative of a good fit). This was achieved for all runs except for one backward run (gap = 9.68%) using GDC-0068 input data. For each network run, we have provided the COSMOS script and its output as separate text files, including all run-specific settings and final gap values (https://osf.io/tdvur/).

Subsequent network analysis and visualization was performed in R, using the rCy3^64^ package to interface with Cytoscape^65^. For the final visualization, a filter was applied such that text was only displayed for nodes with betweenness values of ≥ 0.05, the size of the text is indicative of the degree, and the color of the node indicative of its COSMOS-derived activation value. Betweenness is a measure of the number of shortest paths going through a node, i.e. how much a node acts as point of connection or information transmission^32^.

### Cancer Cell Line Growth Inhibition Screen

The high throughput cell line screen was outsourced to Horizon by Astra Zeneca. A detailed description of the protocol, alongside cell-line specific culture conditions and GI50 curve fits, are included on the OSF project webpage (https://osf.io/us45v/). Briefly, the 288 cell lines were thawed and expanded until they reached their expected doubling times, at which point the screening begins. Cells were seeded in 25 µl of growth media in black 384-well tissue culture and equilibrated at 37°C and 5% CO_2_ for 24 h before treatment. At the time of treatment, a set of assay plates were collected for initial (V_0) measurements of ATP (used as proxy for viability) using the luminescence-based CellTiter Glo 2.0 (Promega) assay and an Envision plate reader (Perkin Elmer). Compounds were transferred to the remaining treatment plates using an Echo acoustic liquid handling system; 25 nl of each compound was added at the appropriate concentration for all dose points. Plates were incubated with compound for 6 days, followed by ATP measurements with CellTiter Glo. All data points were collected via automated processes and subjected to quality control, followed by analysis using Horizon’s proprietary software.

Horizon utilizes Growth Inhibition (GI) as a measure of cell growth. The GI percentages are calculated by applying the following test and equation:

*If* T<V_0 : 100*(1-(T-V_0)/V_0)

*If* T≥ V_0 : 100*(1-(T-V_0)/(V-V_0))

where T is the signal measure for a test drug, V is the untreated/vehicle-treated control measure, and V_0 is the untreated/vehicle control measure at time zero (see above). This formula is derived from the Growth Inhibition calculation used in the National Cancer Institute’s NCI-60 high throughput screen.

#### Computational Integration with DepMap Datasets

Publicly available transcriptomic, proteomic and RPPA data, alongside the relevant metadata, for breast cancer cells of interest were retrieved from DepMap using the *depmap* R package. PCA, GSEA, hierarchical clustering and heatmap generation as part of subsequent integration with experimental GI50adj data were performed as described for RNA sequencing (see “*Raw read mapping, counting and differential expression*”). RNAseq data were obtained as *transcrips per million* (TPM) and were subject to quantile-based filtering (quantile threshold = 0.25), using the *TCGAbiolinks* package, to remove lowly-expressed genes. We used non-parametric Spearman’s correlation to measure the strength of association between variables of interest. GI50adj values were log-transformed (base10) for visualization. For RPPA data, all antibodies labeled with “Caution” were excluded from analysis; the remaining antibody measurements were converted to z-scores prior to visualization. Annotated scripts for the associated analysis steps are provided on the OSF project webpage (https://osf.io/us45v/).

### Pharmacodynamic Analyses in BT474C Xenografts

BT474C pharmacodynamic animal studies were conducted according to AstraZeneca’s Global Bioethics Policy in accordance with the PREPARE and ARRIVE guidelines. Female Nude mice were surgically implanted with a 0.36 mg/60d 17β-estradiol pellet (Innovative Research of America) into the left subcutaneous flank. The following day BT474C cells were implanted at 5 x 10^6^ cells per mouse (suspended in 50% DMEM:50% Matrigel) into the right subcutaneous flank. Mouse weights were monitored twice weekly up until dosing, after which mouse weights were monitored daily. Tumors were measured twice weekly by caliper, with tumor volumes calculated using the formula:

Volume = (π x Maximum measure(Length or Width) x Minimum measure(Length or Width) x Minimum measure (Length or Width))/6000

Mice were randomized by tumor volume into either control or treatment groups when average tumor volume reached 0.5 cm^3^. GDC-00068 was dosed perorally twice a day for 4 days at 12.5 mg/kg (5 ml/kg)(0.5% HPMC, 0.1% Tween 80). INY-05-040 and INY-03-041 were dosed for 4 days as a once daily intraperitoneal injection at 25mg/kg (5ml/kg) (10% DMSO/20% Captisol, pH 5.0 with gluconic acid). On the final day of dosing, 4 h post dosing AM dose, mice were humanely killed, and tumor tissue was collected and immediately snap frozen in liquid nitrogen before storage at -80 °C.

Protein was extracted from snap-frozen tumor fragments by adding 900 μL of extraction buffer (20 mM Tris at pH7.5, 137 mM NaCl, 10% Glycerol, 50 mM NaF, 1 mM Na3VO4, 1% SDS, 1% NP40 substitute) with complete protease inhibitor cocktail (Roche Cat. #11836145001; 1 tablet per 50 mL). Samples were homogenized twice for 30 seconds at 6.5m/s in a fast-prep machine with an incubation at 4 °C for 5 min between runs. Lysates were then sonicated in a chilled Diagenode Bioruptor for two cycles (setting: HIGH) of 30 s ON/30s OFF. Lysates were cleared twice by centrifugation, followed by protein concentration estimation with the Pierce BCA Protein Assay Kit (Thermo Fisher Scientific Cat. # 23227). Approximately 40 μg of protein was run on a NuPAGE 4–12% Bis-Tris gel (Thermo Fisher Scientific) using standard methods. Following protein separation, protein was transferred onto nitrocellulose membranes using dry transfer with iBlot2 (Thermo Fisher Scientific #IB21001). Primary antibodies were diluted in Tris-buffered saline (TBS)/0.05% Tween (TBS/T) supplemented with 5% Marvel, and incubated overnight at 4 °C. The membranes were washed three times for 15 min each in 20 ml of TBS/T. A secondary horseradish peroxidase (HRP)-linked antibody was diluted 1:2000 in TBS/T supplemented with 5 % Marvel and incubated for 1 h at room temperature. The membranes were washed three times for 15 min each in 20 mL of TBS/T, and the signal detected using chemiluminesent SuperSignal West Dura Extended Duration Substrate (Thermo Fisher Scientific).

### Statistics and Reproducibility

Sample size for individual experiments was not pre-determined. Statistical analyses on multidimensional datasets are detailed in the relevant methods sections. To avoid the pitfalls of significance testing on conventional, low-throughput biological datasets, effect size statistics for data shown in Figure 4 were generated using DABEST (Data Analysis using Bootstrap-Coupled ESTimation)^66^.

The exact number of technical and biological replicates are specified in the relevant figure legends. We use biological replicates to refer to independent experimental repeats or tumor samples from different mice. Technical replicates refer to individual samples exposed to the same treatment within the same experimental replicate. The biological responses deduced from the Western blots shown in Figures 1 and 4 were reproduced across independent biological contexts (Figures S1 and S6, respectively); for all other Western blots performed in T47D cells only, all independent experimental replicates are included in the Supplementary Material as indicated. Raw Western blot images are available on the OSF project webpage: https://osf.io/maq7k/.

## Acknowledgements

We thank members of the Toker laboratory, Joan Brugge (Harvard Medical School), Steve Elledge (Harvard Medical School), Kevin Haigis (Dana Farber Cancer Institute), Taru Muranen (Harvard Medical School), Philip Cole (Harvard Medical School), and Benjamin Turk (Yale School of Medicine), for helpful advice and suggestions; John Asara and Min Yuan for technical support with metabolomics. We would also like to thank Karen Adelman, Seth Goldman, and the Nascent transcriptomics Core (Harvard Medical School) for assistance with PRO-Seq library construction and data analysis.

## Author contributions

A.T. and N.S.G. conceived the concept of AKT degraders and supervised the project. R.R.M. analyzed omics datasets, performed COSMOS, and supervised the project. I.Y. designed and synthesized the compounds, and conducted experiments required for their initial chemical and biological characterization. E.C.E. designed and performed all other cell culture-based experiments. G.P. provided technical assistance. A.D. assisted with the COSMOS installation and subsequent result interpretation. K.A.D. performed proteomics experiments, under the supervision of E.S.F. J.W.J. and S.T.B. contributed to the design of studies and supervised work performed within AstraZeneca. R.E.Z. contributed to the synthesis of INY-05-040. C.C. supervised the cell panel profiling and analyzed the raw data. S.W. and J.I.M. performed xenograft experiments. S.R. performed pharmacodynamics profiling and contributed technical assistance. E.C.E., R.R.M. and A.T. wrote the manuscript and prepared figures for publication. All authors reviewed the final manuscript.

## Data and materials availability

Source data and annotated analysis workflows are available on the following OSF project website: https://osf.io/3ay2w/. Transcriptomic data (RNAseq and PRO-seq) have been deposited with GEO under series accession number GSE206389. Proteomics data have been deposited under PRIDE accession number PXD036614 (the data will become publicly available upon manuscript publication). Any outstanding information regarding the computational work will be provided by the corresponding author, Dr Ralitsa Madsen, upon request. A complete list of reagents is provided in Tables 1-4. All custom reagents will be provided by the corresponding author, Dr Alex Toker, upon request.

## KEY RESOURCES TABLES

**Table 1:** Primary Antibodies.

| 1° antibody | Mol.<br>Weight<br>(kDa) | Lot # | Vendor | Cat. # |
| --- | --- | --- | --- | --- |
| pan-AKT | 60 | 20 | Cell Signaling Technology | 4691 |
| pan-AKT | 60 |  | Cell Signaling Technology | 9275 |
| pAKT (S473) | 60 |  | Cell Signaling Technology | 4060 |
| pPRAS40 (T246) | 40 | 12 | Cell Signaling Technology | 2997 |
| pPRAS40 | 40 |  | Cell Signaling Technology | 13175 |
| Total PRAS40 | 40 | 11 | Cell Signaling Technology | 2691 |
| Total PRAS40 | 40 |  | Cell Signaling Technology | 2610 |
| pGSK3β (S9) | 46 | 13 | Cell Signaling Technology | 9336 |
| GSK3β | 46 | 14 | Cell Signaling Technology | 9315 |
| pTSC2 (T1462) | 200 | 7 | Cell Signaling Technology | 3617 |
| TSC2 | 200 | 2 | Cell Signaling Technology | 3990 |
| Vinculin | 124 | 6 | Cell Signaling Technology | 13901 |
| Vinculin | 124 |  | Abcam | AB13007 |
| p-p44/42 (ERK1/2)<br>(T204/202) | 42, 44 | 12 | Cell Signaling Technology | 9101 |
| ERK1/2 | 42, 44 | 21 | Cell Signaling Technology | 4695 |
| 4EBP | 15-20 | 10 | Cell Signaling Technology | 9452 |
| p-S6 (S240/244) | 32 | 7 | Cell Signaling Technology | 5364 |
| p-S6 (S235/236) | 32 |  | Cell Signaling Technology | 4858 |
| S6 | 32 | 9 | Cell Signaling Technology | 2217 |
| p-p38 MAPK (T180/Y182) | 43 | 10 | Cell Signaling Technology | 4511 |
| p38 MAPK | 40 | 9 | Cell Signaling Technology | 8690 |
| p-cJun (S73) | 48 | 5 | Cell Signaling Technology | 3270 |
| cJun | 48 | 13 | Cell Signaling Technology | 9165 |
| pAKT (S473) | 60 | 24 | Cell Signaling Technology | 4060 |
| pAKT (T308) | 80 | 18 | Cell Signaling Technology | 2965 |

**Table 2:** Secondary Antibodies.

| 2° Antibody | Vendor | Lot # | Cat. # | Dilution |
| --- | --- | --- | --- | --- |
| IRDye 800CW Goat anti-Rabbit IgG (H + L) | LI-COR | D00825-14 | 926–32211 | 1:20,000 |
| IRDye® 680LT Goat anti-Mouse IgG (H + L), 0.5 mg | LI-COR | D01014-04 | 926–68070 | 1:20,000 |
| Anti-rabbit IgG, HRP-linked | CST |  | 7074 | 1:2000 |
| Anti-mouse IgG, HRP-linked | CST |  | 7076 | 1:2000 |

**Table 3:** Chemicals & Reagents.

| Reagent | Source | Cat. # |
| --- | --- | --- |
| RPMI-1640 | Wisent Bioproducts | 350000CL |
| DMSO | Fisher Scientific | BP231-100 |
| Bortezomib (PS-241) | Cayman Chemical | 10008822 |
| Pevonedistat (MLN4924) | Selleckchem | S7109 |
| JNK-IN-8 | MedChemExpress | HY-13319 |
| MK-2206 | Cayman Chemical | 11593 |
| Borussertib | Selleckchem | S8839 |
| VH032 | MedChemExpress | HY-120217 |
| AZD 5363 | Cayman Chemical | 15406 |
| Triton X-100 | Fisher | BP151-100 |
| CellTox Green Cytotoxicity Assay | Promega | G8743 |
| NucleoSpin RNA Plus | Takara | 740984.50 |
| Nitrocellulose membrane | Bio Rad | 1620112 |
| Milk | Fisher | NC9022655/190915ASC |
| Bovine Serum Albumin (BSA) | Goldbio | A-421-10 |
| DC Protein Assay Reagent A | Bio Rad | 500-0113 |
| DC Protein Assay Reagent B | Bio Rad | 500-0114 |
| TBS-T | Boston Bioproducts | IBB-181-4L |
| TBS | Boston Bioproducts | IBB_596 |
| SDS Running Buffer | Boston Bioproducts | BP-150 |
| Transfer Buffer | Boston Bioproducts | BP-190 |
| Methanol | Pharmco | 33900HPLC |
| Fetal Bovine Serum | GeminiBio | A020003 |
| Cell Titer Glo | Promega | G7570 |
| Tandem Mass Tag (TMT) Reagents | Thermo Fisher Scientific | A34808 |
| cOmplete, Mini Protease Inhibitor Cocktail | Roche | 11836153001 |
| PhosSTOP, Phosphatase Inhibitor Tablets | Roche | 4906837001 |
| BCA Protein Assay Kit | ThermoFisher | 23227 |
| Protease Inhibitor Cocktail | Roche | 11836145001 |
| Nucleospin RNA Plus Kit | Takara | 740984.50 |
| Nitrocellulose membrane | BioRad | 1620112 |
| PageRuler Plus | Fisher | PI26619 |
| Protease Inhibitor Cocktail | Sigma-Aldrich | P8340-5ML |
| Triton X-100 | Fisher Scientific | BP151-100 |
| 96-well TC treated plates | ThermoFisher | 165305 |
| 60 mm TC treated dishes | Corning | 430166 |
| 10 cm TC treated dishes | Westnet | 353003 |
| 6-well treated tissue culture plates | Greiner | TCG-657160 |
| Black-walled clear-bottom 96-well plates | Fisher | 12-566-70 |
| Mycoplasma Detection Kit | Lonza | LT07-218 |
| 15 cm TC treated dishes | Corning | 08-772-24 |
| NP40 substitute | Roche | 11754599001 |

**Table 4:**
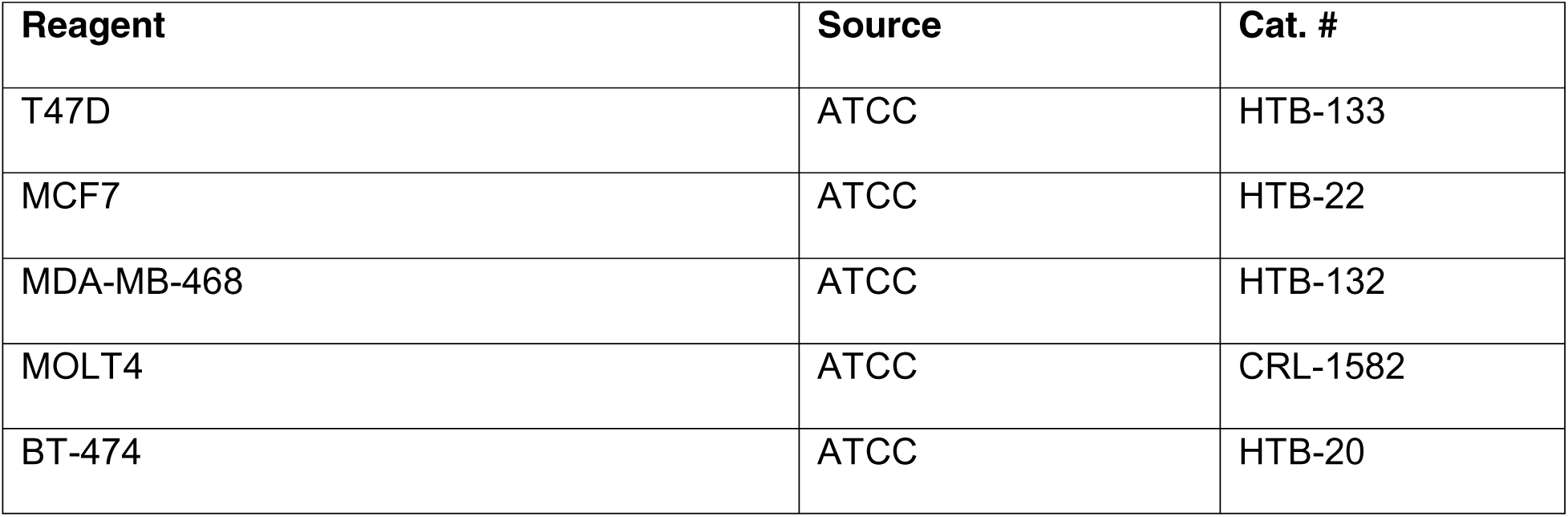
Cell Lines.

**Table 5:** Software & Algorithms.

| Software | Source |
| --- | --- |
| GraphPad Prism | www.graphpad.com/ |
| R Framework | www.R-project.org/ |
| RStudio | https://www.rstudio.com/ |
| Proteome | Thermo Fisher Scientific |
| Discoverer 2.2 |  |
| Adobe Illustrator | www.adobe.com/creativecloud.html |
| Affinity Designer | https://affinity.serif.com/en-gb/designer/ |
| ImageStudioLite | https://www.licor.com/bio/image-studio-lite/ |

## Supplementary Material

Supplementary file with Figures S1-S8.

Additional source data and annotated analysis workflows are available via a bespoke OSF project website (https://osf.io/3ay2w/) with the following subcomponents:

- RNAseq and metabolomics raw and processed data, including annotated analysis scripts (direct access: https://osf.io/3f2m5/)
- PRO-seq raw and processed data, including annotated analysis scripts (direct access: https://osf.io/3f2m5/)
- COSMOS scripts, including source data for all network outputs and tables with run- specific settings (https://osf.io/tdvur/)
- Raw and processed data, including annotated analysis scripts, for CellToxGreen-based cytotoxicity assays (https://osf.io/fasqp/)
- Cell line screen raw and processed data, alongside detailed protocol information, cell line-specific culture conditions and individual GI50 curve fits (https://osf.io/us45v/)
- Raw Western blot images (https://osf.io/maq7k/)

## Figure legends for Supplementary Figures

**Figure S1. INY-05-040 biochemical selectivity and proteomics.** (**A**) TREEspot visualization of the biochemical selectivity profile of GDC-0068 and INY-05-040 (1 μM). AKT isoforms are highlighted in blue, all other inhibited kinases are highlighted in red. (**B**) Scatterplot plot of relative protein abundance changes following 4-h treatment of MOLT4 cells with INY-05-040 (250 nM) *versus* DMSO (vehicle), measured using tandem mass tag quantitative mass spectrometry. The log2 fold-change (FC) is shown on the y axis and - log10(p-value) on the x axis for one independent biological replicate for drug treatment and 3 independent biological replicates for DMSO treatments; *p*-values were adjusted for multiple comparisons (FDR ≤ 0.05). (**C**) Immunoblots for pan-AKT, phospho-PRAS40 (T246), total PRAS40, phospho-S6 (S240/244), total S6, and Vinculin after 5-h treatment of T47D cells with DMSO, INY-05-040 (040) or INY-05-040-Neg at the indicated doses. (**D**) Immunoblots for the same components as in (C), representing MDA-MB-468 cells treated for 5 h with DMSO, INY-05-040 or GDC-0068 at the indicated concentrations. (**E**) Immunoblots for the same components as in (C), representing MDA-MB-468 cells treated with DMSO, INY-05-040 (100 nM) or INY-03-041 (100 nM) for the indicated times. (**F**) Immunoblots for the same components as in (C), representing 5 h co-treatment of T47D and MDA-MB-468 cells with DMSO, bortezomib (0.5 μM), MLN- 4924 (1 μM), and either INY-05-040 (100 nM) or DMSO. (**G**) Immunoblots for the same components as in (C), representing MDA-MB-468 cells treated with DMSO, INY-05-040 (100 nM) or GDC-0068 (100 nM) for 5 h, followed by washout for the indicated times. (**H**) CellTiter Glo assay evaluating percent inhibition in cell growth relative to DMSO treatment in T47D, MCF7, BT-474, or MDA-MB-468 cells, treated for 72 h with INY-03-041, INY-05-040 or GDC-0068. (**I**) Table representing cell line-specific EC50 values (nM) calculated from the respective CellTiter Glo assays in (H).

**Figure S2. Signaling immunoblots related to RNAseq.** Immunoblots for pan-AKT, phospho-PRAS40 (T246), total PRAS40, phospho-GSK3β (S9), total GSK3β, phospho-S6 (S240/244), total S6, and Vinculin after treatment of T47D cells for 5 h or 10 h with DMSO, 100 nM INY-05-040 (040), 100 nM INY-05-040- Neg (Neg), or 500 nM GDC-0068 (GDC), collected in parallel with the corresponding RNAseq samples. Quantification of AKT represents protein abundance over Vinculin, relative to the corresponding DMSO condition for each time point. Quantification of remaining phosphorylated proteins represent normalization to the corresponding total protein, relative to the DMSO signal for each time point.

**Figure S3. Supporting multi-omic data analyses (T47D breast cancer cells), related toFig. 2. (A) + (B)** UpSet intersection plots for up- (A) and downregulated (B) transcripts, respectively, for the indicated treatments relative to DMSO. Fold-change cut-off for differential expression was 1.3; FDR ≤ 0.05. Only genes with HGNC (HUGO Gene Nomenclature Committee) annotation were included in the final count. (**C**) Volcano plot of SREBF1 and SREBF2 target gene expression in Degrader- and GDC-0068-treated T47D cells. The horizontal dotted line indicates the adjusted p-value cut-off for statistical significance (FDR ≤ 0.05); the vertical dotted lines specify the cut-off corresponding to a fold-change of log2(1.3) for up- or downregulation. The target genes correspond to those used for transcription factor footprint estimates with DoRothEA. Black rectangles are used to highlight cholesterol synthesis genes that are selectively upregulated in Degrader- but not GDC-0068-treated cells after 10 h. (**D**) Principal component analysis (PCA) of the PRO-seq dataset, comprising n = 2 independent experiments per treatment (all performed for 5 h). The first three independent axes (principal components; PCs) of highest variation are shown. (**E**) + (**F**) As for (A) and (B), respectively, but using differentially expressed genes from the PRO-seq dataset.

**Figure S4. Signaling immunoblots related to metabolomics.** Immunoblots for pan-AKT, phospho- PRAS40 (T246), total PRAS40, phospho-S6 (S240/244), total S6, and Vinculin after treatment of T47D cells for 24 h with DMSO, INY-05-040, INY-05-040-Neg, GDC-0068, AZD 5363, or MK-2206 as indicated; samples were collected in parallel with the corresponding metabolomics samples. Note that the dose of GDC-0068 was increased to 750 nM in Trial 3 to retain consistent levels of signaling suppression relative to the previous experiments. Quantification of AKT represents protein abundance over Vinculin, relative to the average of the replicate DMSO samples. Quantification of remaining phosphorylated proteins represent normalization to the corresponding total protein, relative to the average of the replicate DMSO samples.

**Figure S5. Individual COSMOS networks following integration of T47D transcriptomic and metabolomic data, specific to Degrader (A) and GDC-0068 (B) treatments.** For details of the analytical framework, refer to Fig. 3A. Predicted inhibitory (-1) and activating (1) interactions are indicated. Predicted average node activity (AvgAct) in each network model is visualized on a scale from -1 (inhibited) to 1 (activated). Each network was generated following an independent COSMOS run with the same data but with varying settings to ensure robustness of the final output (for additional information on run-specific settings, see https://osf.io/tdvur/).

**Figure S6. Stress MAPK signaling activation in MCF7 and MDA-MB-468 cells.** Immunoblots for pan-AKT, phospho-PRAS40 (T246), total PRAS40, phospho-p38α (T180/Y182), total p38α, phospho-c-Jun (S73), total cJun, phospho-S6 (S240/244), total S6, and Vinculin after DMSO, INY-05-040 (100 nM) or GDC-0068 (750 nM) treatment of (**A**) MCF7 or (**B**) MDA-MB-468 cells for the indicated times. Quantification of AKT and cJun represents protein abundance over Vinculin, relative to the corresponding DMSO condition for each time point. Quantification of remaining phosphorylated proteins represent normalization to the corresponding total protein, relative to the DMSO signal for each time point.

**Figure S7. Cell viability after pre-treatment of cells with JNK-IN-8.** (**A**) Cytotoxicity index assayed using CellTox Green, in BT-474 or T47D cells treated for 24 h with either DMSO or the indicated concentrations of JNK-IN-8, followed by 120-h co-treatment with either DMSO, INY-05-040 (100 nM) or GDC-0068 (750 nM). The cytotoxicity index represents cytotoxicity values corrected for background fluorescence and normalized to total signal following chemical permeabilization (used as proxy measure for total cell number). (**B**) Immunoblots for pan-AKT, phospho-p38α (T180/Y182), total p38α, phospho-c-Jun (S73), total cJun, phospho-S6 (S240/244), total S6, PARP (FL: full lengths; CL: cleaved), Vinculin, and beta-actin after 24 h pre-treatment of BT474 cells with either DMSO or 50 nM JNK-IN-8, followed by 120-h co-treatment with either DMSO, INY-05-040 (100 nM) or GDC-0068 (750 nM). Treatment with Bortezomib (10 µM) for 24 h and 48 h was used as positive control. Two technical replicates (indicated with Plate 1 and Plate 2) were processed in parallel. Complementary brightfield microscopy images for both (A) and (B) are provided on the OSF project website (https://osf.io/fasqp/). Quantification for cleaved (CL) PARP was performed by measuring the intensity of the indicated lower band, normalized to beta-actin, relative to DMSO.

**Figure S8. Cancer cell line screen of GDC-0068, INY-03-041, and INY-05-040.** (**A**) Heatmap of cell line-specific GI50adj values for each compound, with Euclidean distance-based clustering of the cell lines (rows). (**B**) Barplots indicating the GI50adj values for each compound in breast cancer cell lines only, colored according to sensitivity to the respective compound (sensitive: GI50adj < 0.5 µM; intermediate: 0.5 µM < GI50adj < 1 µM; resistant: GI50adj > 1 µM). The dotted horizontal line indicates GI50adj = 1 µM.

