## Supplementary Material for "Multi-omic profiling of breast cancer cells uncovers stress MAPK-associated sensitivity to AKT degradation"

This PDF file includes:

Figures S1 to S8

Additional source data and annotated analysis workflows are available via a bespoke OSF project website (<https://osf.io/3ay2w/>) with the following subcomponents:

- RNAseq and metabolomics raw and processed data, including annotated analysis scripts (direct access: <https://osf.io/3f2m5/>)
- PROseq raw and processed data, including annotated analysis scripts (direct access: <https://osf.io/3f2m5/>)
- COSMOS scripts, including source data for all network outputs and tables with run-specific settings (<https://osf.io/tdvur/>)
- Raw and processed data, including annotated analysis scripts, for CellToxGreen-based cytotoxicity assays (<https://osf.io/fasqp/>)
- Cell line screen raw and processed data, alongside detailed protocol information, cell line-specific culture conditions and individual GI50 curve fits (<https://osf.io/us45v/>)
- Raw Western blot images (<https://osf.io/maq7k/>)

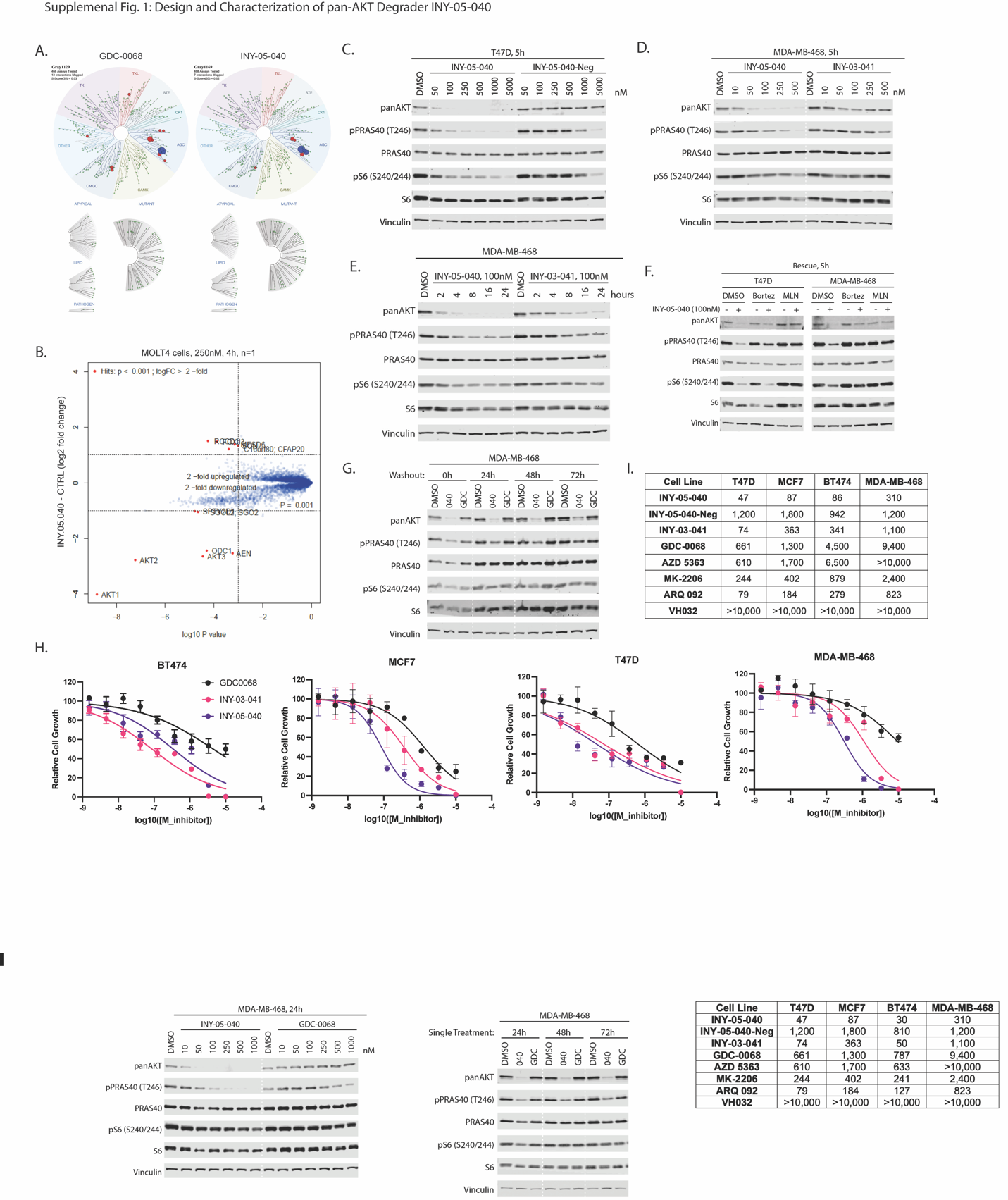

**Figure S1. INY-05-040 biochemical selectivity and proteomics.** (**A**) TREEspot visualization of the biochemical selectivity profile of GDC-0068 and INY-05-040 (1 μM). AKT isoforms are highlighted in blue, all other inhibited kinases are highlighted in red. (**B**) Scatterplot plot of relative protein abundance changes following 4-h treatment of MOLT4 cells with INY-05-040 (250 nM) *versus* DMSO (vehicle), measured using tandem mass tag quantitative mass spectrometry. The log2 fold-change (FC) is shown on the y axis and -log10(p-value) on the x axis for one independent biological replicate for drug treatment and 3 independent biological replicates for DMSO treatments; *p*-values were adjusted for multiple comparisons (FDR ≤ 0.05). (**C**) Immunoblots for pan-AKT, phospho-PRAS40 (T246), total PRAS40, phospho-S6 (S240/244), total S6, and Vinculin after 5-h treatment of T47D cells with DMSO, INY-05-040 (040) or INY-05-040-Neg at the indicated doses. (**D**) Immunoblots for the same components as in (C), representing MDA-MB-468 cells treated for 5 h with DMSO, INY-05-040 or GDC-0068 at the indicated concentrations. (**E**) Immunoblots for the same components as in (C), representing MDA-MB-468 cells treated with DMSO, INY-05-040 (100 nM) or INY-03-041 (100 nM) for the indicated times. (**F**) Immunoblots for the same components as in (C), representing 5 h co-treatment of T47D and MDA-MB-468 cells with DMSO, bortezomib (0.5 mM), MLN-4924 (1 mM), and either INY-05-040 (100 nM) or DMSO. (**G**) Immunoblots for the same components as in (C), representing MDA-MB-468 cells treated with DMSO, INY-05-040 (100 nM) or GDC-0068 (100 nM) for 5 h, followed by washout for the indicated times. (**H**) CellTiter Glo assay evaluating percent inhibition in cell growth relative to DMSO treatment in T47D, MCF7, BT-474, or MDA-MB-468 cells, treated for 72 h with INY-03-041, INY-05-040 or GDC-0068. (**I**) Table representing cell line-specific EC50 values (nM) calculated from the respective CellTiter Glo assays in (H).

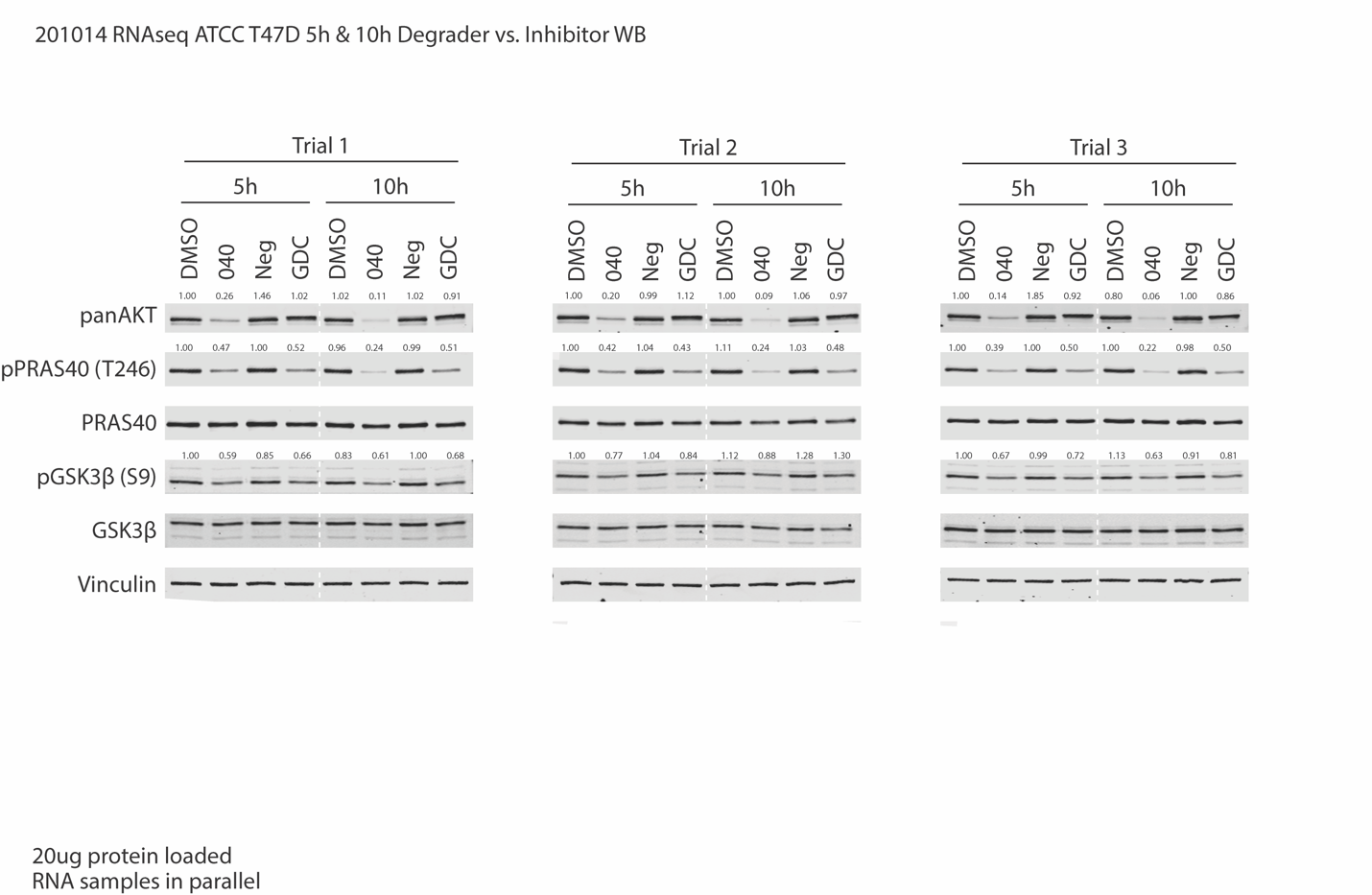

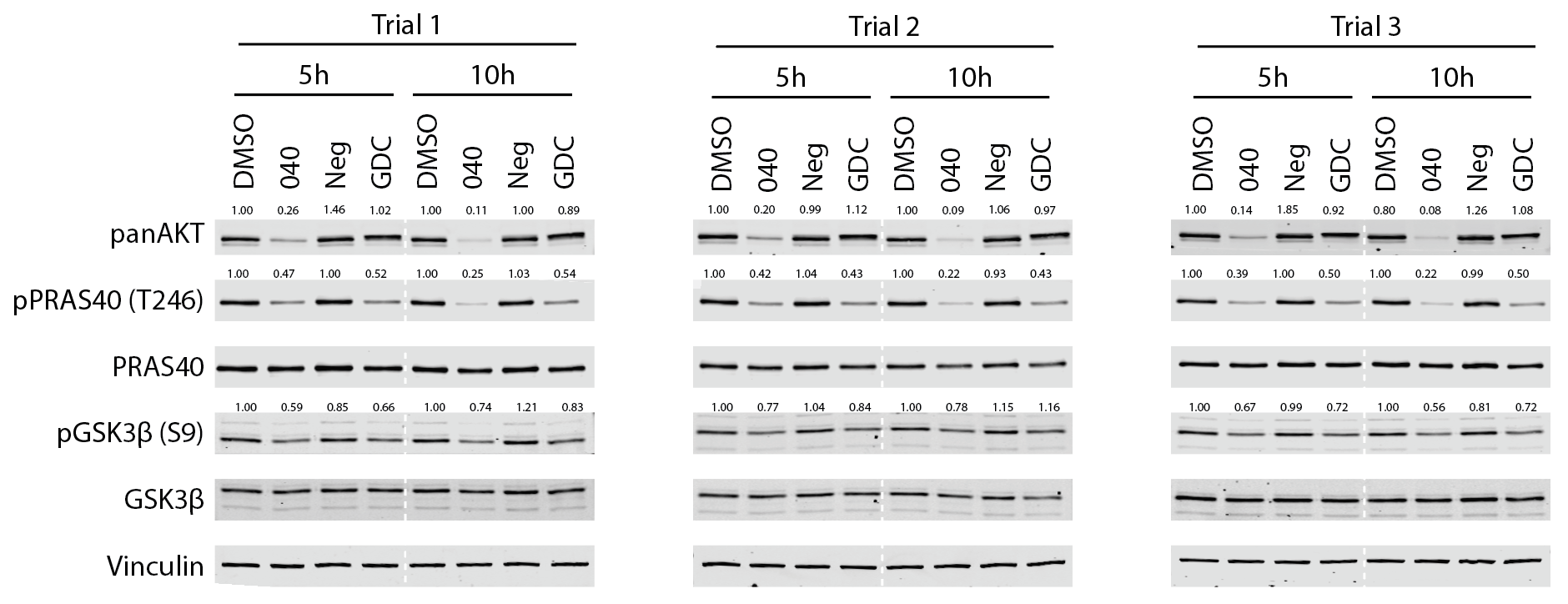

**Figure S2. Signaling immunoblots related to RNAseq.** Immunoblots for pan-AKT, phospho-PRAS40 (T246), total PRAS40, phospho-GSK3β (S9), total GSK3β, phospho-S6 (S240/244), total S6, and Vinculin after treatment of T47D cells for 5 h or 10 h with DMSO, 100 nM INY-05-040 (040), 100 nM INY-05-040-Neg (Neg), or 500 nM GDC-0068 (GDC), collected in parallel with the corresponding RNAseq samples. Quantification of AKT represents protein abundance over Vinculin, relative to the corresponding DMSO condition for each time point. Quantification of remaining phosphorylated proteins represent normalization to the corresponding total protein, relative to the DMSO signal for each time point.

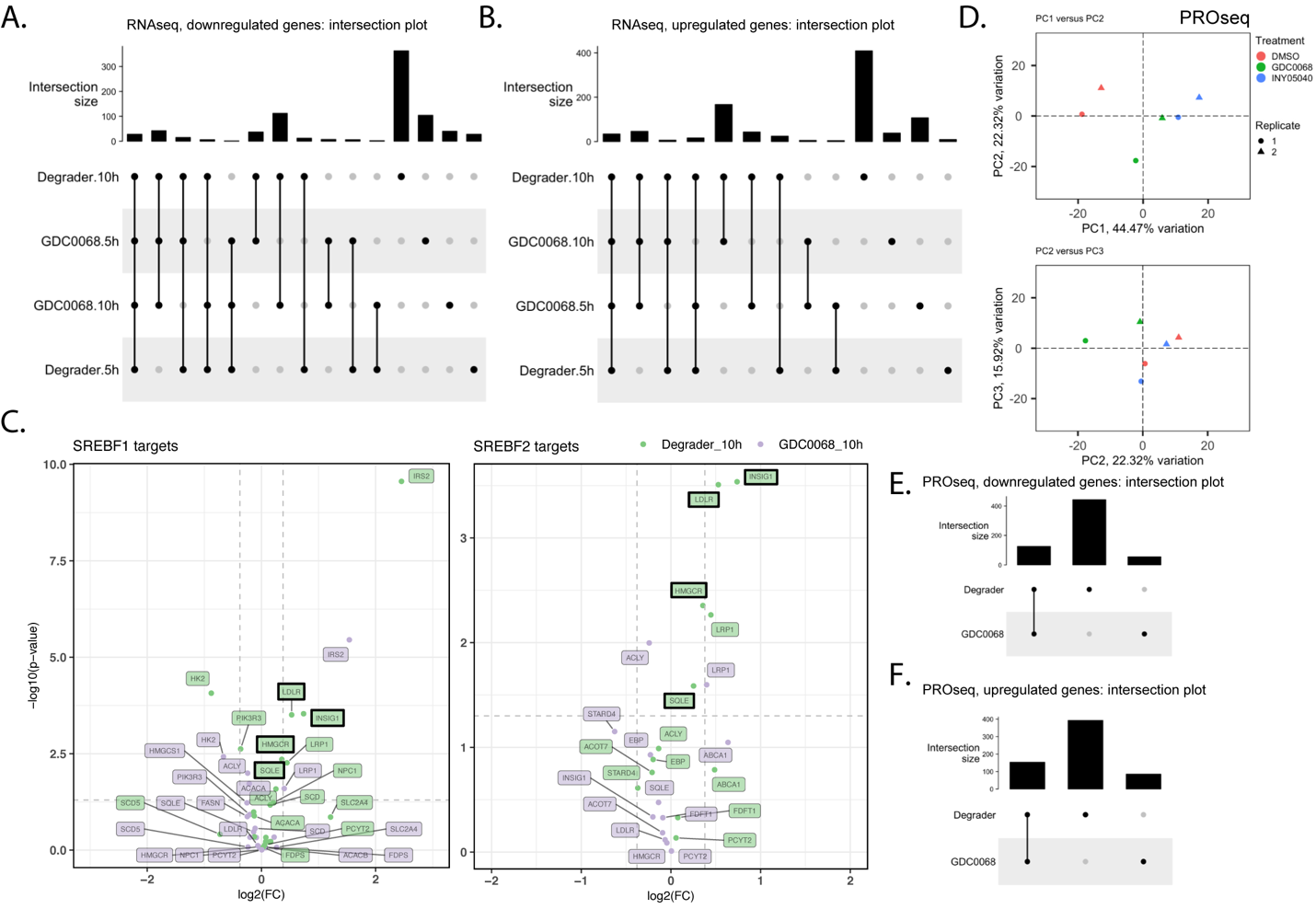

**Figure S3. Supporting multi-omic data analyses (T47D breast cancer cells), related to Fig. 2. (A) + (B)** UpSet intersection plots for up- (A) and downregulated (B) transcripts, respectively, for the indicated treatments relative to DMSO. Fold-change cut-off for differential expression was 1.3; FDR ≤ 0.05. Only genes with HGNC (HUGO Gene Nomenclature Committee) annotation were included in the final count. (**C**) Volcano plot of SREBF1 and SREBF2 target gene expression in Degrader- and GDC-0068-treated T47D cells. The horizontal dotted line indicates the adjusted p-value cut-off for statistical significance (FDR ≤ 0.05); the vertical dotted lines specify the cut-off corresponding to a fold-change of log2(1.3) for up- or downregulation. The target genes correspond to those used for transcription factor footprint estimates with DoRothEA. Black rectangles are used to highlight cholesterol synthesis genes that are selectively upregulated in Degrader- but not GDC-0068-treated cells after 10 hours. (**D**) Principal component analysis (PCA) of the PROseq dataset, comprising n = 2 independent experiments per treatment (all performed for 5 h). The first three independent axes (principal components; PCs) of highest variation are shown. (**E**) + (**F**) As for (A) and (B), respectively, but using differentially expressed genes from the PROseq dataset.

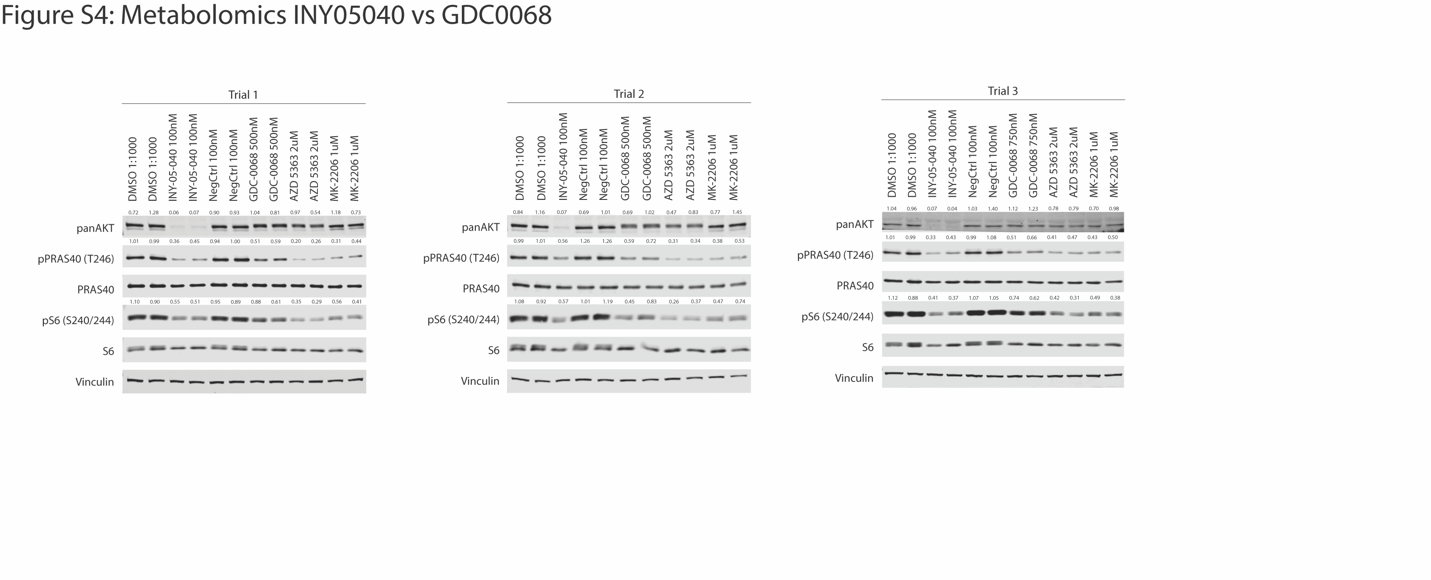

**Figure S4. Signaling immunoblots related to metabolomics.** Immunoblots for pan-AKT, phospho-PRAS40 (T246), total PRAS40, phospho-S6 (S240/244), total S6, and Vinculin after treatment of T47D cells for 24 h with DMSO, INY-05-040, INY-05-040-Neg, GDC-0068, AZD 5363, or MK-2206 as indicated; samples were collected in parallel with the corresponding metabolomics samples. Note that the dose of GDC-0068 was increased to 750 nM in Trial 3 to retain consistent levels of signaling suppression relative to the previous experiments. Quantification of AKT represents protein abundance over Vinculin, relative to the average of the replicate DMSO samples. Quantification of remaining phosphorylated proteins represent normalization to the corresponding total protein, relative to the average of the replicate DMSO samples.

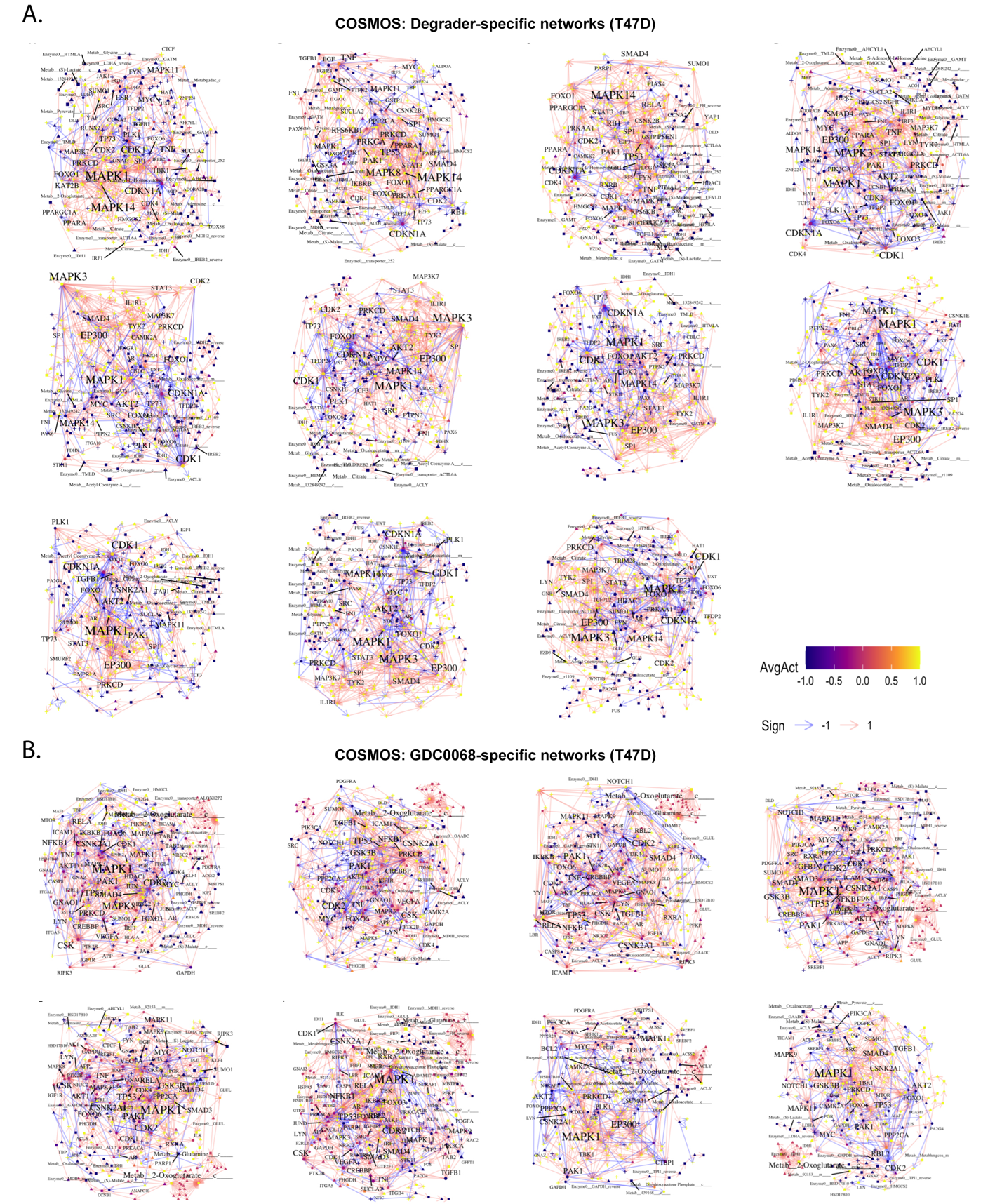

**Figure S5. Individual COSMOS networks following integration of T47D transcriptomic and metabolomic data, specific to Degrader (A) and GDC-0068 (B) treatments.** For details of the analytical framework, refer to Fig. 3A. Predicted inhibitory (-1) and activating (1) interactions are indicated. Predicted average node activity (AvgAct) in each network model is visualized on a scale from -1 (inhibited) to 1 (activated). Each network was generated following an independent COSMOS run with the same data but with varying settings to ensure robustness of the final output (for additional information on run-specific settings, see <https://osf.io/tdvur/>).

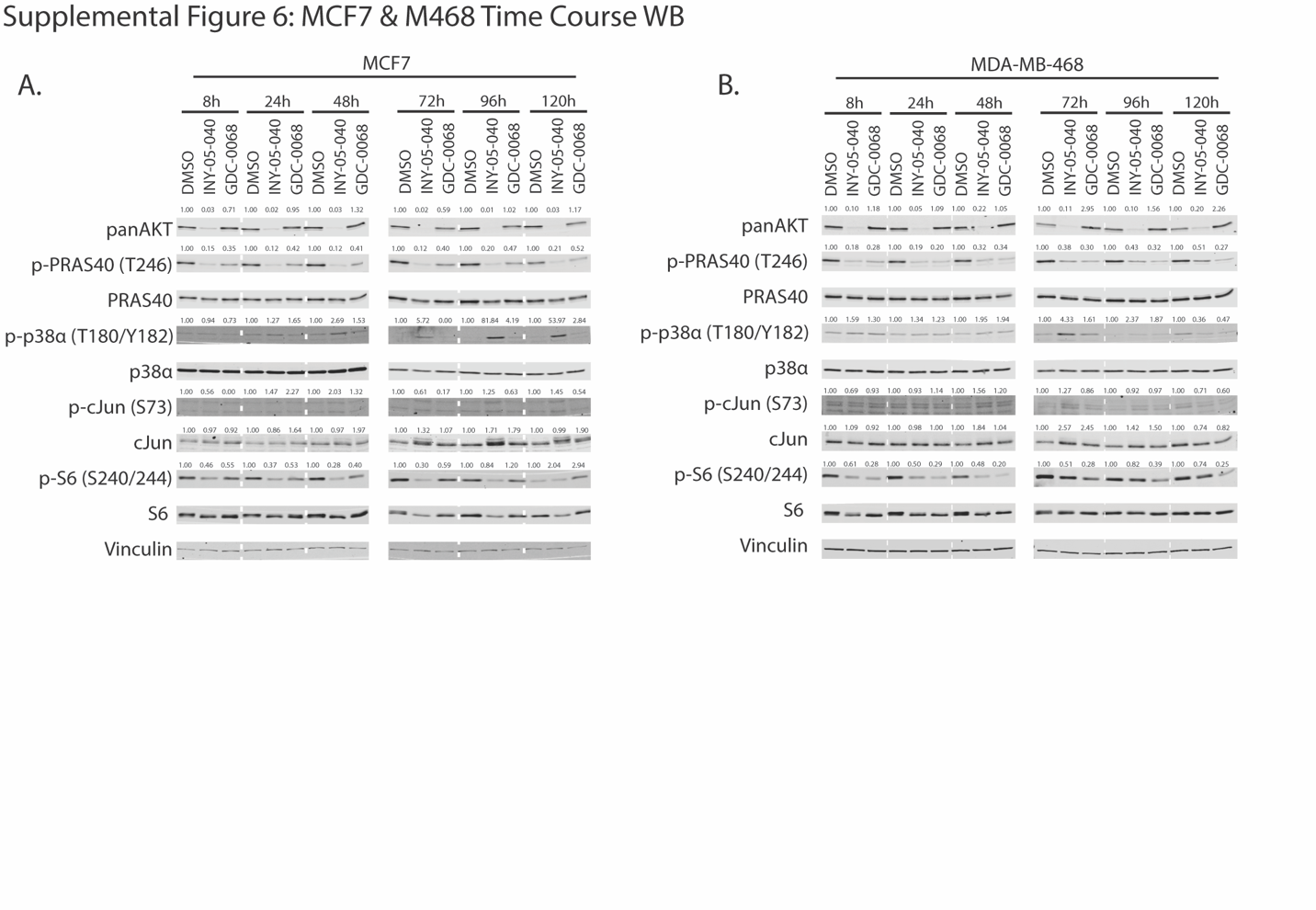

**Figure S6. Stress MAPK signaling activation in MCF7 and MDA-MB-468 cells.** Immunoblots for pan-AKT, phospho-PRAS40 (T246), total PRAS40, phospho-p38α (T180/Y182), total p38α, phospho-c-Jun (S73), total cJun, phospho-S6 (S240/244), total S6, and Vinculin after DMSO, INY-05-040 (100 nM) or GDC-0068 (750 nM) treatment of (**A**) MCF7 or (**B**) MDA-MB-468 cells for the indicated times. Quantification of AKT and cJun represents protein abundance over Vinculin, relative to the corresponding DMSO condition for each time point. Quantification of remaining phosphorylated proteins represent normalization to the corresponding total protein, relative to the DMSO signal for each time point.

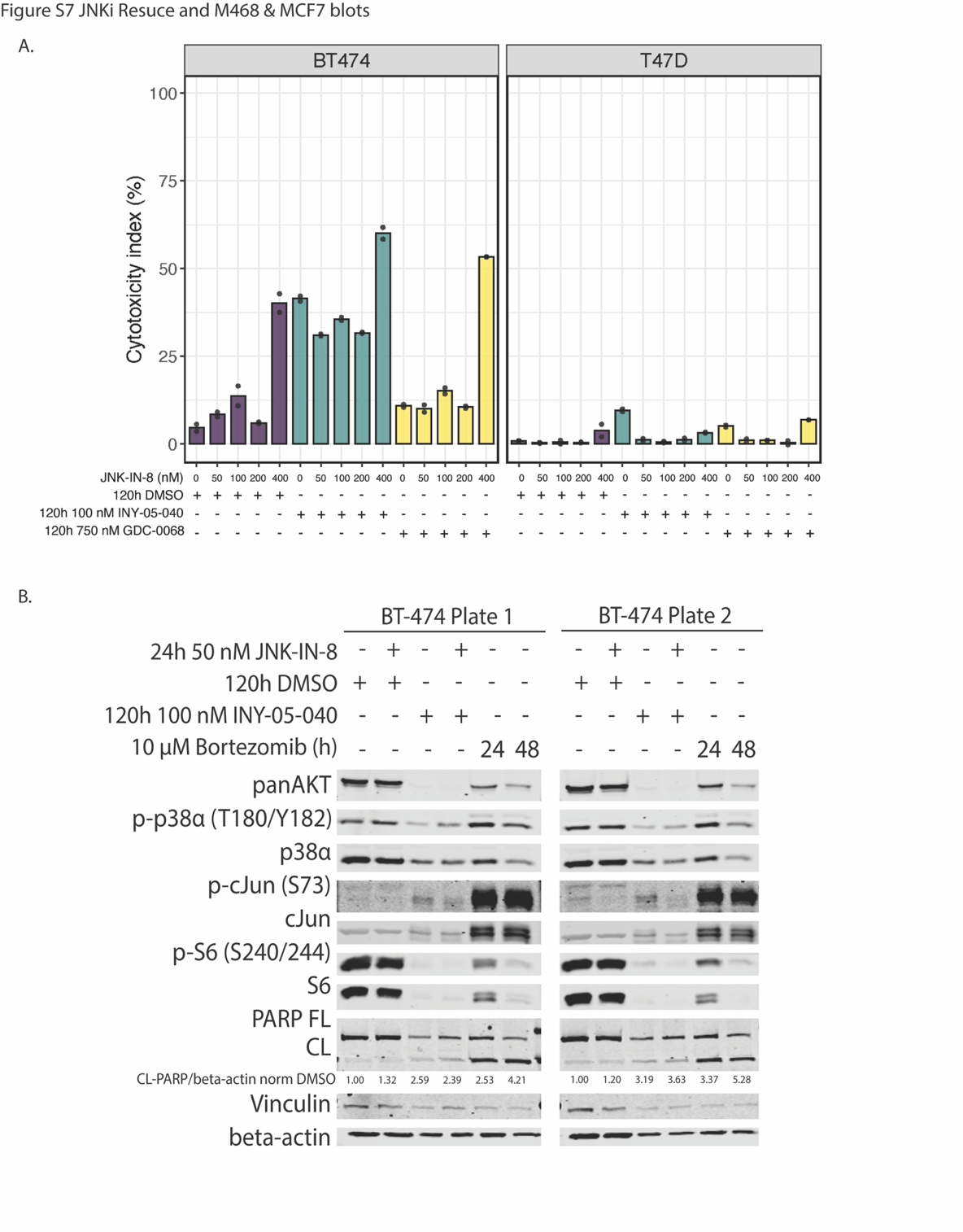

**Figure S7. Cell viability after pre-treatment of cells with JNK-IN-8.** (**A**) Cytotoxicity index, assayed using CellTox Green, in BT-474 or T47D cells treated for 24 h with either DMSO or the indicated concentrations of JNK-IN-8, followed by 120-h co-treatment with either DMSO, INY-05-040 (100 nM) or GDC-0068 (750 nM). The cytotoxicity index represents cytotoxicity values corrected for background fluorescence and normalized to total signal following chemical permeabilization (used as proxy measure for total cell number). (**B**) Immunoblots for pan-AKT, phospho-p38α (T180/Y182), total p38α, phospho-c-Jun (S73), total cJun, phospho-S6 (S240/244), total S6, PARP (FL: full lengths; CL: cleaved), Vinculin, and beta-actin after 24 h pre-treatment of BT474 cells with either DMSO or 50 nM JNK-IN-8, followed by 120-h co-treatment with either DMSO, INY-05-040 (100 nM) or GDC-0068 (750 nM). Treatment with Bortezomib (10 µM) for 24 h and 48 h was used as positive control. Two technical replicates (indicated with Plate 1 and Plate 2) were processed in parallel. Complementary brightfield microscopy images for both (A) and (B) are provided on the OSF project website (<https://osf.io/fasqp/>). Quantification for cleaved (CL) PARP was performed by measuring the intensity of the indicated lower band, normalized to beta-actin, relative to DMSO.

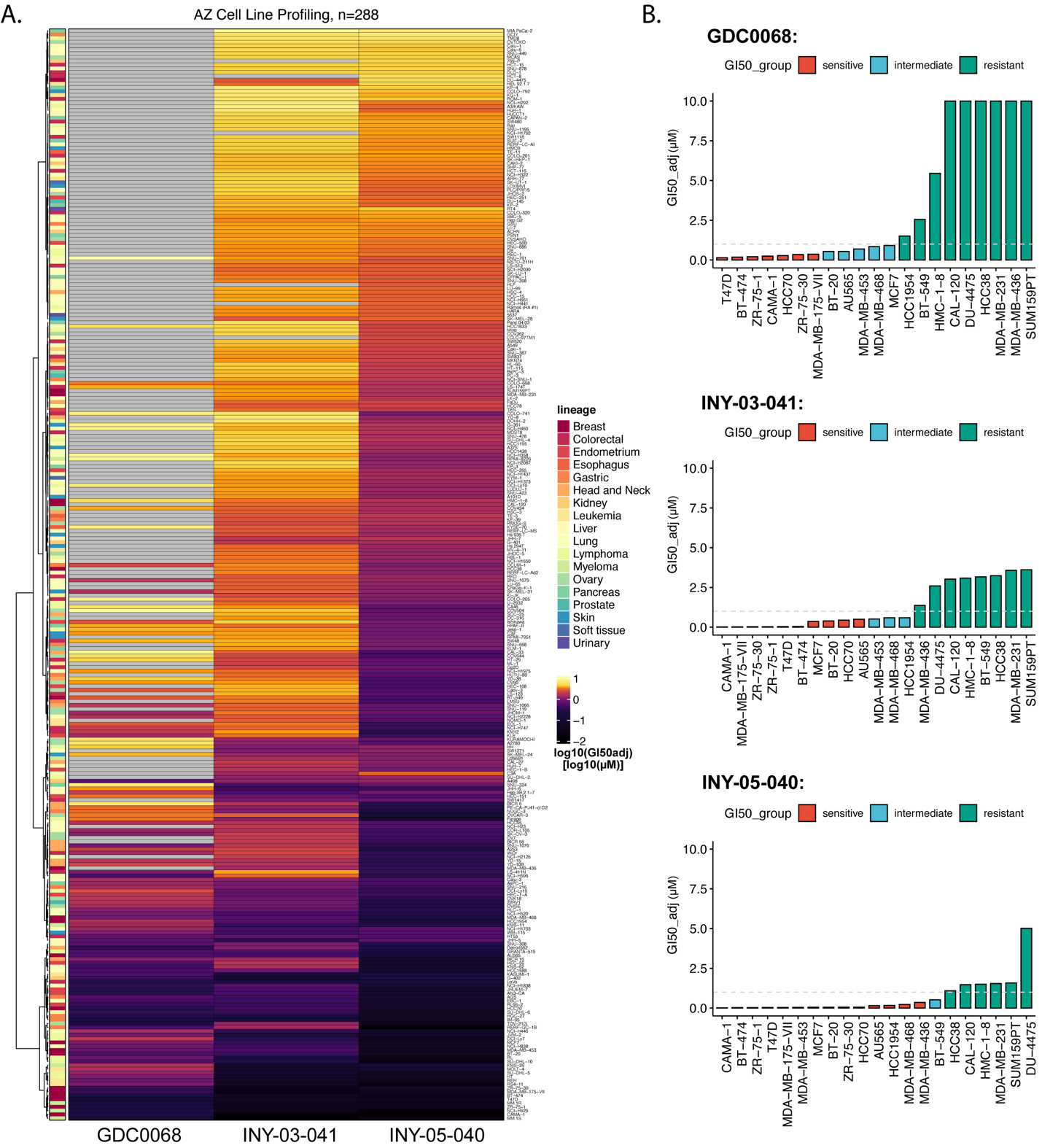

**Figure S8. Cancer cell line screen of GDC-0068, INY-03-041, and INY-05-040.** (**A**) Heatmap of cell line-specific GI50adj values for each compound, with Euclidean distance-based clustering of the cell lines (rows). (**B**) Barplots indicating the GI50adj values for each compound in breast cancer cell lines only, colored according to sensitivity to the respective compound (sensitive: GI50adj < 0.5 µM; intermediate: 0.5 µM < GI50adj < 1 µM; resistant: GI50adj > 1 µM). The dotted horizontal line indicates GI50adj = 1 µM.
